# Accurate and fast graph-based pangenome annotation and clustering with ggCaller

**DOI:** 10.1101/2023.01.24.524926

**Authors:** Samuel T. Horsfield, Nicholas J. Croucher, John A. Lees

**Author notes:** Contributed equally (Nicholas J. Croucher, John A. Lees). Author emails: - Samuel T. Horsfield - Nicholas J. Croucher - John A. Lees.

## Abstract

Bacterial genomes differ in both gene content and sequence mutations, which can cause important clinical phenotypic differences such as vaccine escape or antimicrobial resistance. To identify and quantify important variants, all genes within a population must be predicted, functionally annotated and clustered, representing the ‘pangenome’. Despite the volume of genome data available, gene prediction and annotation are currently conducted in isolation on individual genomes, which is computationally inefficient and frequently inconsistent across genomes. Here, we introduce the open-source software graph-gene-caller (ggCaller; https://github.com/samhorsfield96/ggCaller). ggCaller combines gene prediction, functional annotation and clustering into a single step using population-wide de Bruijn Graphs, removing redundancy in gene annotation, and resulting in more accurate gene predictions and orthologue clustering. We applied ggCaller to simulated and real-world bacterial genome datasets, comparing it to current state-of-the-art tools. ggCaller is ~50x faster with equivalent or greater accuracy, particularly in datasets with complex sources of error, such as assembly contamination or fragmentation. ggCaller is also an important extension to bacterial genome-wide association studies, enabling querying of annotated graphs for functional analyses. We highlight this application by functionally annotating DNA sequences with significant associations to tetracycline and macrolide resistance in *Streptococcus pneumoniae*, identifying key resistance determinants that were missed when using only a single reference genome. ggCaller is a novel bacterial genome analysis tool with applications in bacterial epidemiology and evolutionary study.

## Introduction

Accurate representation of population’s genomic diversity, known as a pangenome, is critical in epidemiological and evolutionary studies of bacterial species. Identification of core genes, found in all individuals, is used for classification and phylogenetic analyses. Such methods include phylogenetic analysis (Wolf *et al*., 2002; Croucher *et al*., 2013b; Zakham *et al*., 2021), and transmission chain inference during outbreaks (Ypma, van Ballegooijen & Wallinga, 2013; Croucher & Didelot, 2015). Genes present in only a subset of isolates, known as accessory genes, are often correlated with particular strains (Lees *et al*., 2019). These genes have also been associated with the wide phenotypic diversity found in many bacterial species, including antimicrobial resistance (AMR) (Jaillard *et al*., 2017; McNally *et al*., 2019), virulence (Alikhan *et al*., 2018; Hennart *et al*., 2020), host range (Dearlove *et al*., 2015; Weinert *et al*., 2015) and vaccine escape (Lo *et al*., 2019). Accessory genes are the focus of many evolutionary models of bacterial population structure and dynamics, such as understanding how multi-strain populations emerge and are maintained (Baumdicker, Hess & Pfaffelhuber, 2012; Iranzo *et al*., 2019; Harrow *et al*., 2021), and predicting how they respond to perturbations such as vaccines (Corander *et al*., 2017; Azarian *et al*., 2020).

Pangenome studies rely on gene prediction in each isolate genome assembly followed by similarity-based clustering, generating clusters of orthologous genes (COGs). These steps are currently run as separate bioinformatic processes, split into gene prediction tools, or gene-callers, and pangenome analysis tools. Gene-callers, such as Glimmer (Delcher *et al*., 2007), Prodigal (Hyatt *et al*., 2010) and GeneMarkS-2 (Lomsadze *et al*., 2018), predict the locations of coding sequences in individual genomes using models of gene structure and gene overlap penalisation. There has been little recent innovation in gene prediction algorithms; a comprehensive benchmarking study of existing tools included only one tool released in the last 10 years, and highlighted no tool was universally applicable across bacteria (Dimonaco *et al*., 2022). Contrastingly, gene annotation, whereby gene prediction tools are integrated with annotation databases to assign functional labels to predicted genes, has seen increased attention. Popular examples include PGAP (Tatusova *et al*., 2016), Prokka (Seemann, 2014), DFAST (Tanizawa, Fujisawa & Nakamura, 2018) and Bakta (Schwengers *et al*., 2021). As gene prediction and annotation tools are designed for analysing single genomes only, a key issue when applying these tools in pangenome studies is the consistency in prediction and annotations across orthologues. For example, if predicted start or stop positions vary between orthologues (termed a ‘prediction error’), under-clustering can occur, whereby truly homologous genes do not share enough sequence to be placed in the same cluster (Zhou, Charlesworth & Achtman, 2020). Orthologues may also be given inconsistent functional annotations (termed an ‘annotation error’), leading to ambiguity during functional inference of gene families (Tonkin-Hill *et al*., 2020). Moreover, functional annotations are applied to genes individually, generating huge computational redundancy, as orthologues are annotated in each genome, rather than once within the population. This leads to increased runtime, ultimately limiting the size and therefore comprehensiveness of the annotation database that can be used (Schwengers *et al*., 2021). Finally, poor assembly quality, such as contamination and fragmentation, can impact gene prediction accuracy by introducing false positive predictions, such as contaminant genes or partial gene sequences (Tonkin-Hill *et al*., 2020). Gene prediction and annotation, specifically for pangenome studies, require innovations to ensure orthologues are identified consistently across a population.

Pangenome analysis tools cluster the predicted gene sequences from all input genomes, representing the pangenome as a gene presence/absence matrix. In practice, clusters are generated first based on sequence similarity, with paralogs being identified using either synteny-based (Tonkin-Hill *et al*., 2020; Page *et al*., 2015) or tree-based approaches (Zhou, Charlesworth & Achtman, 2020; Ding, Baumdicker & Neher, 2018). Roary (Page *et al*., 2015), developed in the first generation of these tools, generates COGs based on a single BLAST threshold without correction for gene mis-annotation. Later tools introduced lower identity thresholds to better cluster divergent gene families, with additional processing to reduce the effects of gene prediction and annotation errors. Panaroo (Tonkin-Hill *et al*., 2020) uses synteny and population-frequency information to identify spurious COGs originating from contaminants or fragmentation, and predicts genes that may have been missed initially by gene-callers. Panaroo also corrects annotation errors by only keeping the best-supported annotation within a COG. However, as Panaroo relies entirely on the input gene sequences and cannot directly correct them, it is sensitive to gene prediction errors. PEPPAN (Zhou, Charlesworth & Achtman, 2020) addresses this issue by generating COGs initially, before identifying the most frequent sequence for each COG and searching for its homologues within all genomes in the dataset. This process ensures all gene start and stop coordinates within a COG are predicted consistently. However, PEPPAN does not employ the same stringent quality-control methods employed in Panaroo, making it susceptible to errors originating from low quality assemblies. Both Panaroo and PEPPAN also rely on gene prediction and annotation within individual genomes, which is computationally inefficient. There is currently no tool that corrects for poor assembly quality and gene prediction errors and avoids redundancy in gene annotation.

To enable non-redundant, consistent, and accurate gene prediction and annotation across a population, a data structure is required that represents the distribution of genetic variation across many genomes. Pangenome graphs provide a means of compacting large collections of linear references into a network where identical or similar sequences are merged into nodes, variation is represented by edges, and individual genomes as paths through the graph (Eizenga *et al*., 2020). De Bruijn graphs are a form of pangenome graph which are built from matching short nucleotide sequences known as k-mers, with edges added between k-mers that share an overlap of *k* – 1 basepairs. Coloured compacted De Bruijn Graphs (from here referred to as DBGs) compress non-branching paths of k-mers into sequences called “unitigs”, with each k-mer being annotated with the genomes, or ‘colours’ in which it is found. DBGs are a highly scalable method of building pangenome graphs, capable of including thousands of bacterial genomes (Holley & Melsted, 2020), and provide a lossless representation of population diversity (Schulz, Wittler & Stoye, 2022). DBGs therefore do not have the redundancy of the equivalent collection of linear genomes, and have the potential to consistently predict and annotate genes, informed by node-level population-frequency. Gene prediction and annotation within a DBG would therefore overcome the issues encountered when conducting pangenome analysis using individual linear genomes.

Here, we present ggCaller (graph gene-caller), a population-wide gene-caller based on DBGs. ggCaller uses population-frequency information to guide gene prediction, aiding the identification of homologous start codons across orthologues, and consistent scoring of orthologues. Moreover, ggCaller removes redundancy from gene annotation, first generating COGs before annotating the entire cluster, instead of each orthologue individually. ggCaller also implements a modified version of Panaroo’s COG graph algorithm to enable removal of spurious COGs and gene refinding. We demonstrate the benefits of DBG-based gene-calling in bacterial datasets using both real and simulated genomes, comparing prediction and annotation accuracy with existing linear genome gene annotation and pangenome analysis tools. We show that ggCaller outperforms state-of-the-art tools in assemblies with sources of error, such as contamination and fragmentation, in terms of recall and precision of COG identification. ggCaller also has greater consistency of gene coordinate prediction when applied to structurally diverse orthologues, whilst significantly reducing runtime. ggCaller can also be applied in pangenome-wide association studies (PGWAS), enabling reference-agnostic functional inference of significant hits. ggCaller is freely available at https://github.com/samhorsfield96/ggCaller, under the open-source MIT licence, and on bioconda.

## Results

### Overview of the ggCaller workflow

ggCaller predicts genes within a DBG, using sequence-sharing across the whole population to guide annotation and clustering of orthologous genes (Figure 1). DBGs are generated by Bifrost (Holley & Melsted, 2020) and can be built either from assemblies or read files in FASTA or FASTQ format (**Step 1**). DBGs are constructed by first matching k-mers (sequences of length k, which is chosen *a priori* by the user). Non-branching paths of k-mers are merged into unitigs (from here referred to as ‘nodes’), compacting the DBG. These nodes are ‘coloured’ based on the input genomes they are found in, enabling calculation of the population frequency of all sequences > *k* bases in length.

**Figure 1:**
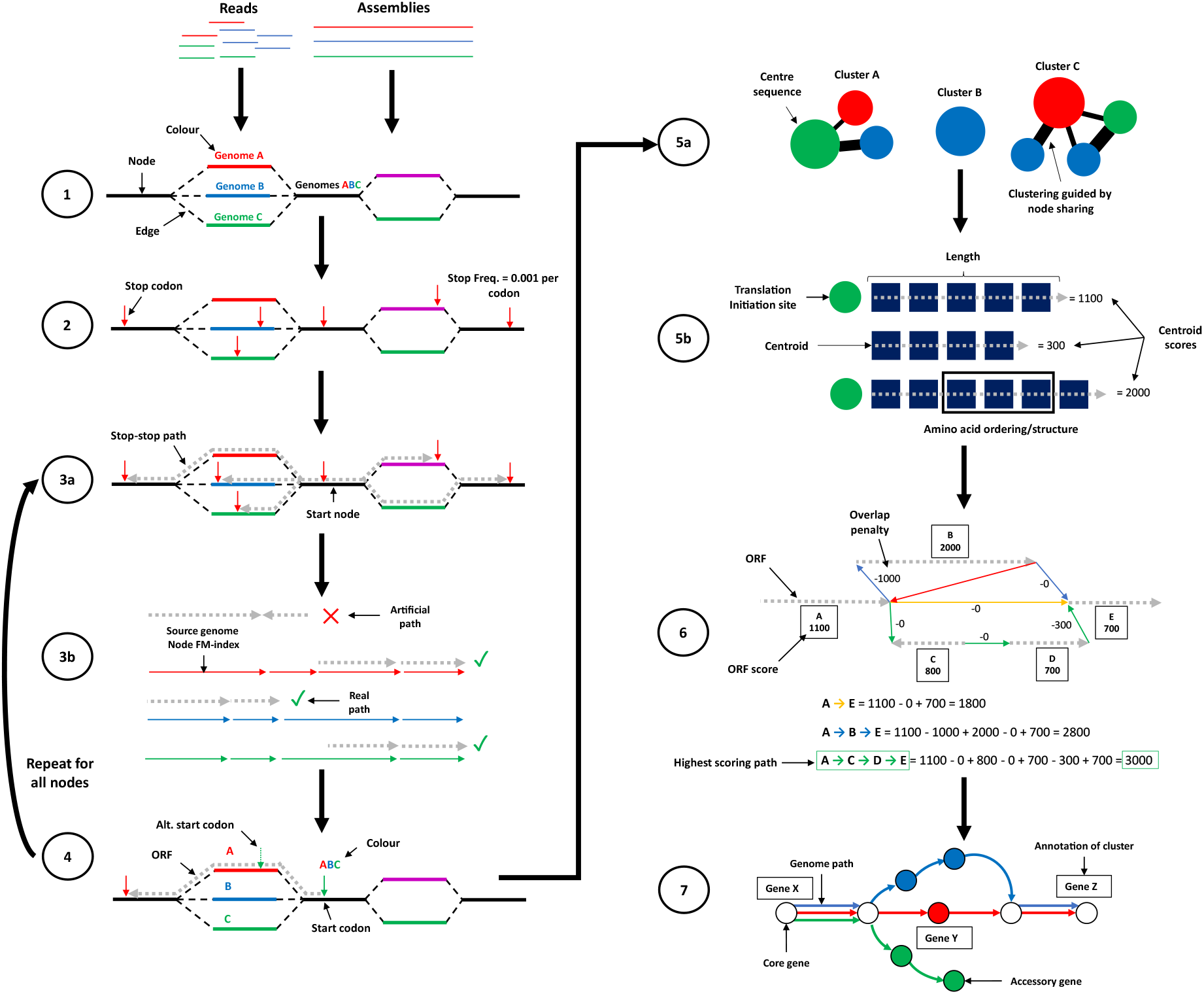
ggCaller workflow. **1)** DBG is generated from reads or assemblies by Bifrost. **2)** All stop codons are identified and stop frequency is calculated (total number of stop codons in DBG / total number of codons in DBG). **3a)** Starting at an initial node containing a stop codon, a depth first search (DFS) is applied to pair all stop codons in the start node with a downstream stop codon in the same reading frame. **3b)** During DFS, paths are compared to an FM-index to remove incorrect paths. **4)** ORFs are defined by identifying start codons scored based on translation initiation site sequence, genome coverage (given by number of colours shared in node) and frequency of this start being chosen in other potential orthologues. Steps **3** and **4** are repeated for all nodes containing a stop codon. **5a)** ORFs are clustered into COGs, using node-sharing to reduce search space. **5b)** BALROG is used to generate an average per-residue score using only the centre sequence of each COG. To score each ORF in the cluster, the per-residue score is multiplied by the respective ORF length. **6)** Highest scoring tiling path calculated for overlapping genes within the DBG using Bellman-Ford algorithm, producing ‘true’ gene-call set. **7)** Gene-calls and synteny information are used to build a COG graph. A modified version of Panaroo is used to remove poorly supported gene-calls, annotate clusters and re-call missed genes/pseudogenes.

ggCaller then identifies all stop codons in the DBG (**Step 2**) and traverses the DBG to identify ORFs (**Step 3**). Each stop codon is paired with a downstream stop-codon in the same reading frame using a depth-first search (**Step 3a**), thereby delineating the coordinates of all possible reading frames. Paths in DBGs do not necessarily represent real contiguous sequences from input genomes (Břinda, Baym & Kucherov, 2021), because repeats within a single genome can link distant sequences in the graph. To remove these artefacts, candidate paths are searched against the contiguous input sequences using an FM-index rank query (**Step 3b**).

The sequences between pairs of stop codons are then searched for start codons, which are paired with downstream stop codons in the same reading frame, generating an open-reading frame (ORF). As bacterial genes have alternative start sites due to reuse of start codons within an exon (Dimonaco *et al*., 2021), the best supported start site is chosen based on three criteria (**Step 4**). For each potential start site, a score is calculated from the start site population-frequency, a translation initiation site (TIS) score using a sequence-scoring model from BALROG (Sommer & Salzberg, 2021), and how many times the start site has been ‘chosen’ before as the best supported start in other potential orthologues. The start site with the highest score is chosen as the correct coordinate, generating a single ORF for each pair of stop codons.

ORFs are then clustered using a method based on Linclust (Steinegger & Söding, 2018) to avoid conducting all-by-all pairwise comparisons. ORFs that share at least one node are placed in a ‘node-group’, with ORFs being compared only to the longest sequence (termed ‘centre’) of each node-group they belong to (**Step 5a**). As each ORF will likely traverse multiple nodes, it will belong to multiple node-groups. Within each node-group, Edlib (Šošić & Šikić, 2017) is used to rapidly calculate pairwise edit distances in amino-acid space, with ORFs being clustered with the centre sequence with which they share the highest identity. Sequences are then scored using the gene sequence-scoring model in BALROG; a temporal convolutional network that generates an average per-residue score for a translated ORF sequence (**Step 5b**). Only centre sequences are scored, vastly reducing the number of BALROG model queries required to score all ORFs. The centre sequence average per-residue score is then multiplied by the respective lengths of each ORF in the cluster, generating an individual score for all ORFs. ORF scores are then used to determine the highest-scoring tiling path through the DBG per input genome, which penalises large overlaps between adjacent ORFs (**Step 6**). This generates a population-wide set of coding sequence (CDS) predictions.

CDS predictions are then passed to an updated version of Panaroo’s COG graph algorithm (Tonkin-Hill *et al*., 2020) that has been adapted to work directly with DBGs rather than linear genomes. COGs are clustered further down to 50% identity, paralogous gene clusters are split, and poorly supported clusters are removed (**Step 7**). This step generates a graph with nodes representing COGs, rather than DNA sequences as used in the DBG. ggCaller uses the same three pre-sets for COG pruning as implemented in Panaroo: sensitive, moderate, and strict (See **Methods**). Clusters are also functionally annotated using DIAMOND (Buchfink, Xie & Huson, 2014) and/or HMMER3 (Eddy, 2009). As in **Step 5b**, only cluster centre sequences are queried, with the functional annotation being shared across all CDSs in the cluster. ggCaller also implements a DBG-based gene refinding module, which enables re-calling of genes or pseudogenes missed on the first pass by ggCaller. The final default outputs are a gene presence/absence matrix, a set of annotated gene clusters and their respective sequences, and their locations in their respective linear input sequences (as standardised GFF3 files). Additionally, core/pangenome alignments, phylogenies and single nucleotide polymorphism calls can be generated automatically.

ggCaller features several innovations over existing gene annotation and pangenome analysis tools. **Steps 2-4** ensure start positions of orthologues are called consistently across a population by considering population frequencies of start codons. This process was implemented to avoid incorrect ORF truncation or extension, which is an issue with one-by-one linear genome gene-calling. **Step 5** reduces the number of BALROG model queries, and ensures orthologues are scored equally. Only scores for cluster centre sequences are generated, which can then be shared across orthologues which have the same or similar scores due to sequence similarity. Similarly in **Step 7**, ggCaller functionally annotates clusters using only centre sequences during orthologue clustering. Both processes were designed to reduce annotation inconsistency and redundancy across orthologs to lower runtime and increase gene prediction and annotation accuracy.

### ggCaller accurately predicts genes from simulated draft assemblies of structurally diverse operons

To initially benchmark gene-annotation accuracy from ggCaller, we predicted genes in a collection of five pneumococcal capsular polysaccharide biosynthetic operons (*cps*) (Bentley *et al*., 2006), comparing predictions with the previously annotated gene coordinates. These *cps* operons were chosen as they are highly diverse in sequence content and structure, consisting of between 16-23 genes, and are manually curated, providing an ideal initial ground-truth dataset. Larger population genome datasets are infeasible to manually curate, and so would lack a ground-truth gene set for benchmarking. To simulate the assembly errors seen in draft assemblies, analysis was conducted on fully intact *cps* sequences, and sequences where all annotated genes were synthetically fragmented with a single contig break at a random position. Genes were predicted with ggCaller in moderate mode, GeneMarkS-2 and Prokka (which uses Prodigal for gene prediction). Pangenome analysis for gene predictions from GeneMarkS-2 and Prokka was conducted using Panaroo. We compared tools based on their recall and precision of ground-truth gene set, and the length of these sequences covered by the respective predictions for each gene-caller to determine how predictions were affected by fragmentation.

Recall and precision of exact matches to ground-truth gene sequences were compared across the original and fragmented *cps* operons (**Figure 2A**). ggCaller performed similarly to other tools in terms of recall in the original *cps* operons, however had slightly lower precision. This can be attributed to differences in gene-structure models, which also explains variation between Prokka and GeneMarkS-2. However, in fragmented *cps* operons, ggCaller was the only tool able to recall any genes with correct start and end coordinates. Tools were also compared based on the proportion of the curated gene length covered by their respective gene-calls (**Figure 2B**). In the original *cps* operons, most gene predictions from all tools fully covered their respective ground-truth sequence. However, in the fragmented *cps* operons, ground-truth sequences were covered to a greater degree by ggCaller predictions (median=0.70) than with Prokka or GeneMarkS-2 (both tools had median=0.55). As DBGs connect k-mers with a *k* – 1 overlap anywhere in the population (Holley & Melsted, 2020), contig breaks in individual assemblies can be spanned by forming a path across k-mers in other assemblies which do not have contig breaks at that orthologous position. This enables ggCaller to recall a greater number of full gene sequences in highly fragmented assemblies than linear genome gene-callers.

**Figure 2:**
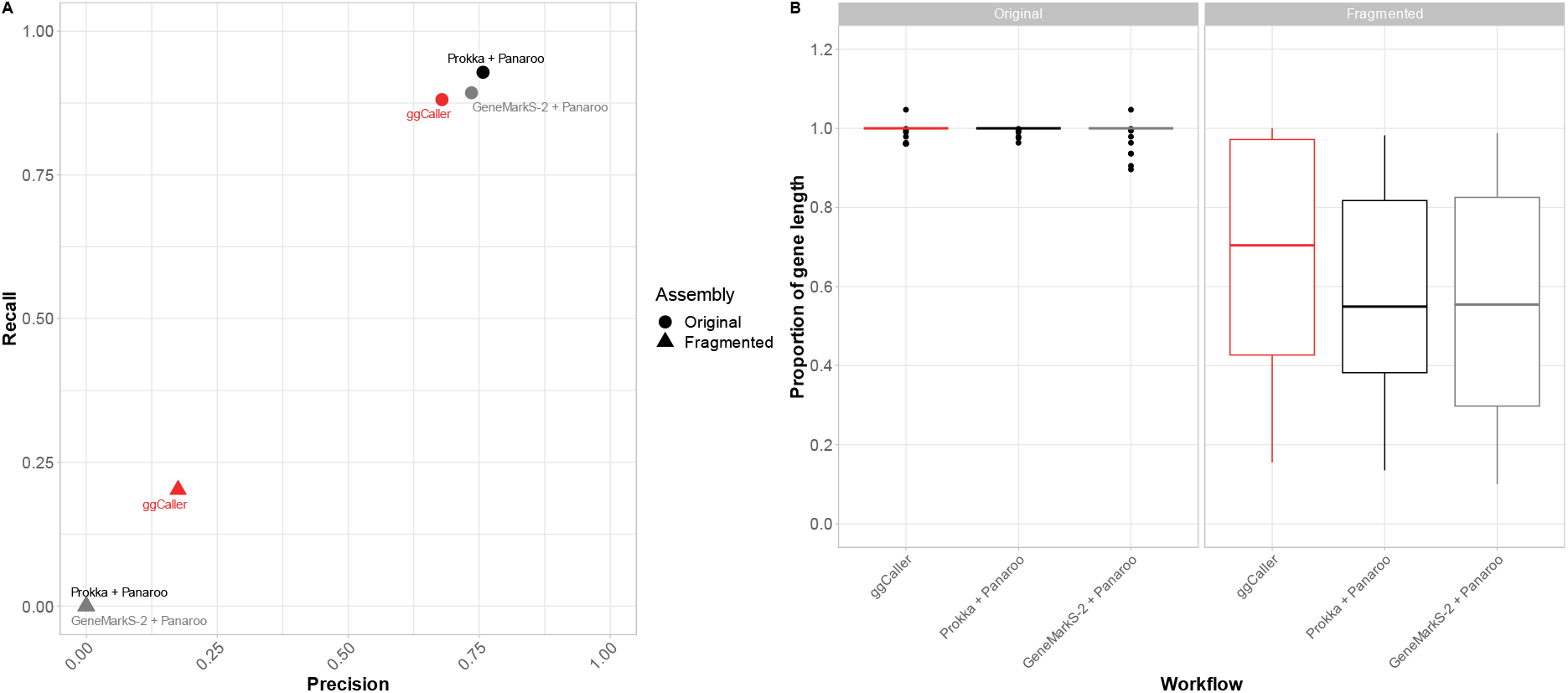
Gene prediction comparison for pneumococcal capsular biosynthetic operons. Precision vs. recall comparisons for correctly identified genes (i.e. correct start and end coordinates) (**A**) and proportion of ground-truth gene length if 3’ end is correctly called (**B**) for original and fragmented ground-truth genes within *cps* operons.

### ggCaller has superior performance in simulated datasets with complex sources of assembly error

In addition to gene prediction, ggCaller provides a single workflow for annotation, orthologue clustering and pangenome analysis. To benchmark ggCaller against existing pangenome analysis workflows, we generated simulated populations of 100 assemblies starting from *Streptococcus pneumoniae* ATCC 700669 serotype 23F (also referred to as ‘Spn23F’) using the workflow described in Tonkin-Hill *et al*., (2020). Briefly, we simulated populations containing 100 genomes, using varying gene gain/loss ratios and within-gene mutation rates, as well as additional fragmentation or contamination. This resulted in seven separate parameter combinations. To more accurately simulate the real-world processes involved in pangenome analysis, assemblies were generated from simulated genomes using ART to generate error-prone reads, and SPADES to assemble these (Huang *et al*., 2012; Bankevich *et al*., 2012). ggCaller was compared against three workflows: genes were first identified and annotated by Prokka, and pangenome analysis was conducted either using either Roary, Panaroo or PEPPAN. We then compared the workflows based on their estimates of total pangenome size and core genome size (defined as COGs present in >99% of genomes) compared to the expected number of COGs provided by the simulation to determine pangenome representation accuracy.

Comparisons of estimated total pangenome and core genome sizes are shown in Figure 3. Simulations were split into simple **(Figure 3A**) and complex **(Figure 3B**) based on their respective parameters (results for simulations not shown above can be found in **Supplementary Figures 1** & **2**). Increasing within-gene mutation rate, decreasing gene gain/loss ratio and introducing contamination or fragmentation had notable negative effects on accuracy for all workflows, causing greater deviation from the expected number of COGs. However, for all simulations, ggCaller produced the most accurate estimates core genome size, independent of stringency settings. In the fragmented simulation, ggCaller core genome size estimations differed from expected values by −91, −92 and −110 COGs for sensitive, moderate, and strict modes respectively. As highlighted in Figure 2, ggCaller can recall a greater number of intact genes in highly fragmented assemblies, which improves clustering accuracy by generating fewer truncated orthologues. In contrast, Panaroo, PEPPAN and Roary core genome size estimations differed by −493, −295 and −1160 COGs respectively. These results highlight an issue in Panaroo and Roary when many assemblies in the dataset are highly fragmented-as these tools rely on gene-order to guide clustering, when this is incorrect or inconsistent in the input (Tonkin-Hill *et al*., 2020; Page *et al*., 2015), under-clustering of COGs occurs. PEPPAN was less sensitive to fragmentation due to use of gene-trees in addition to gene-order to generate COGs, reducing the effect of assembly fragmentation (Zhou, Charlesworth & Achtman, 2020), however it was still less accurate than ggCaller.

**Figure 3:**
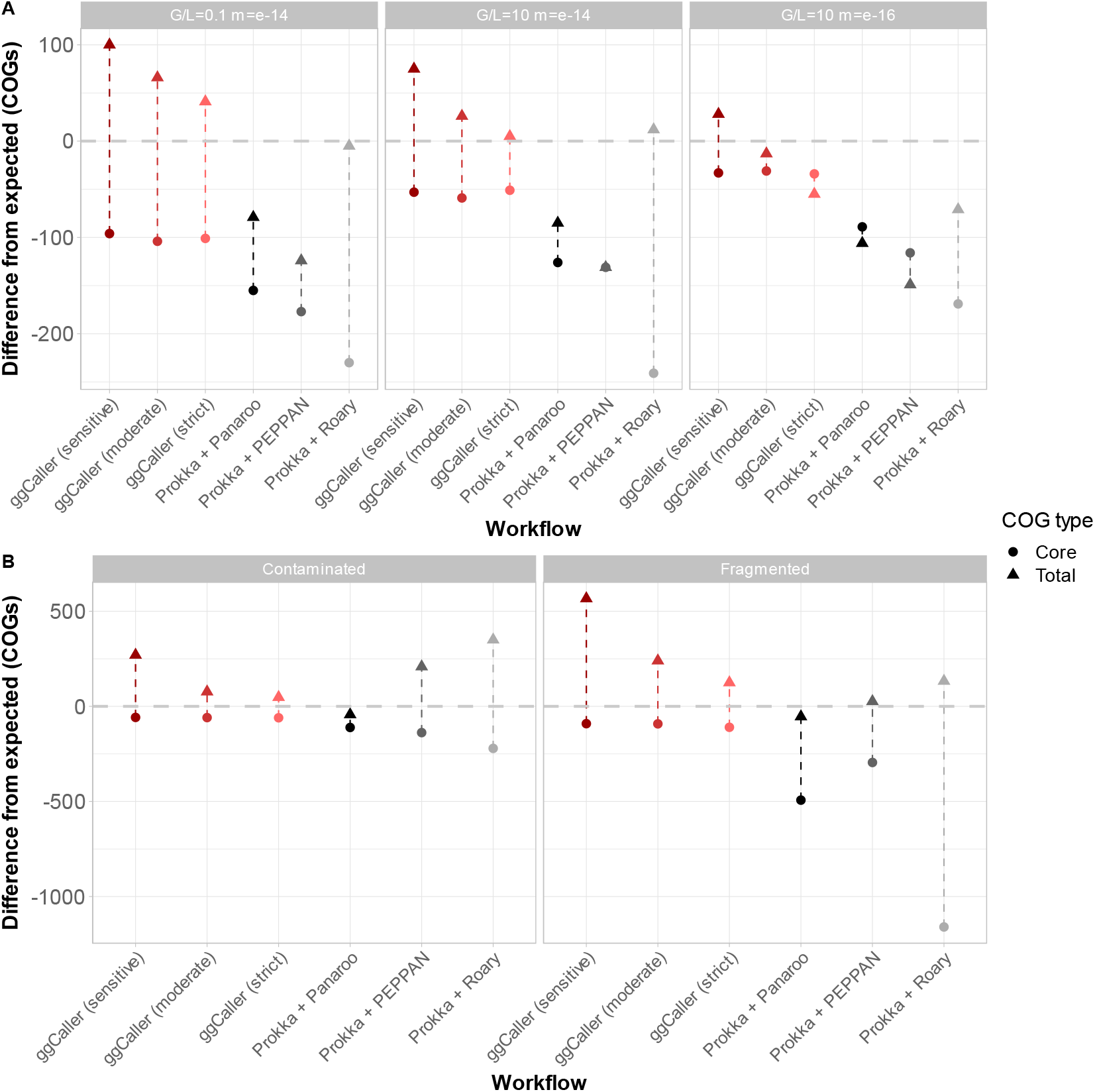
Comparison of estimated total pangenome and core genome sizes across simulated populations. Panels describe simulations with simple (**A**) and complex (**B**) sources of error. Difference from expected refers to the over- or underestimation of the number of COGs from the ground truth in the simulation (grey dotted line). Over- or underestimation will result in a positive or negative difference respectively. If a workflow exactly estimates the core/total pangenome size, there will be no difference from the expected number of COGs. G/L refers to the gene gain to gene loss ratio, m refers to the mutation rate per site per generation. Total expected pangenome size for all simulations was 2259 COGs.

For total pangenome size estimates, ggCaller became more accurate with increasing stringency of the Panaroo error-correcting step. Results from ggCaller in moderate and strict modes produced similar estimates of total pangenome size which were the top two most accurate workflows for all simple simulations. In contrast, Panaroo and PEPPAN underestimated total pangenome size, whilst Roary varied between under- and overestimation. For the fragmented simulation, ggCaller predicted a slightly larger total pangenome size than other tools. This is likely a biproduct of ggCaller identifying genes across contig breaks, which can result in false positives. This effect was minimised by increasing stringency, with strict mode estimating +126 COGs compared to PEPPAN, Panaroo and Roary which were +26, −54 and +133 from the ground-truth respectively. In the simulation of contaminated input genomes, ggCaller (moderate and strict) and Panaroo all accurately estimated total pangenome size (+77, +48 and −43 from expected values respectively), whilst PEPPAN and Roary overestimated it by +290 and +350 COGs respectively. ggCaller and Panaroo both employ the same quality control measures to identify and remove low-frequency COGs that are likely contaminants, which are not used in PEPPAN or Roary, resulting in overestimation of total pangenome size.

The recall and precision of expected COGs were also compared to quantify differences in gene annotation and clustering accuracy in simple **(Figure 4A**) and complex **(Figure 4B**) simulations. ggCaller either performed equivalently to, or outperformed, all other tools in all simulations, except for the fragmented simulation. Notably, ggCaller had the fewest false negatives across all simulations, whilst PEPPAN had the highest. The increased number of false positives for ggCaller in the fragmented simulation aligns with the slight overestimation in total pangenome size seen in **Figure 3B**. However, this effect is minor, with ggCaller having 119 and 122 false positives for moderate and strict modes respectively, compared to 84 for both Panaroo and PEPPAN. PEPPAN and Roary had the highest number of false positives in the contaminated simulation (311 and 330 respectively), whereas ggCaller had the lowest of all tools (82, 81 and 82 for sensitive, moderate and strict). Within correctly predicted COGs, ggCaller had similar numbers of errors to Panaroo and PEPPAN, apart from the simulation with increased gene gain/loss ratio (G/L=10, m=e-14). However, this number was still 25% of that of Roary (**Supplementary Figure 3**). Overall, these simulations show that ggCaller performs as well as, or better than, gold-standard pangenomic analysis workflows in simulated populations, particularly in datasets featuring complex sources of assembly error.

**Figure 4:**
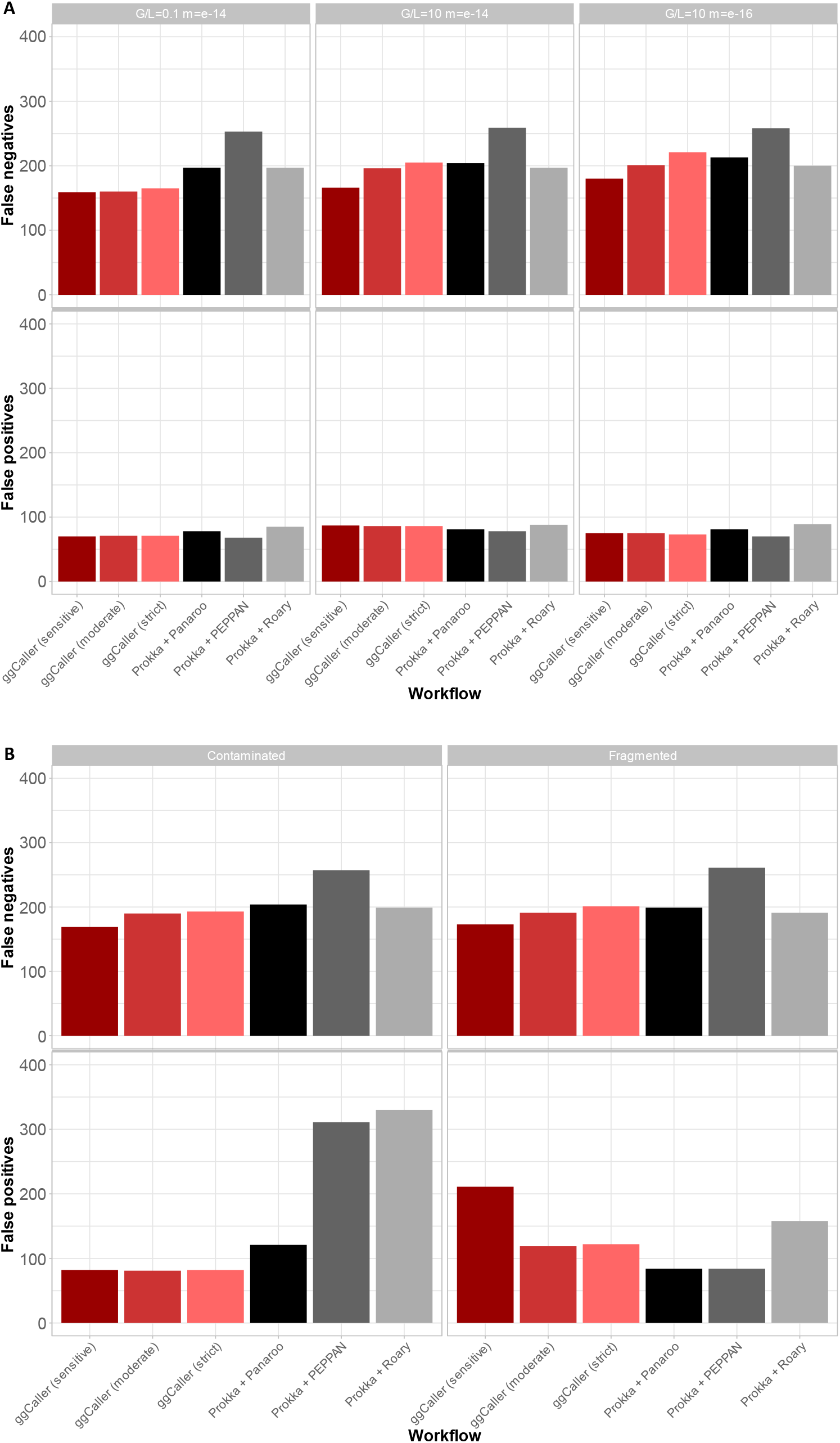
Comparison of COG annotation accuracy across simulated populations. Panels describe simulations withs simple (**A**) and complex (**B**) sources of error. False positives are COGs that were called by a workflow but were not present in the ground-truth set. False negatives are COGs that were present in the ground-truth set but not called by a workflow.

### ggCaller accurately represents pangenomes of bacterial species with varying levels of diversity

To benchmark ggCaller on real-world data, we analysed genome sequences from three bacterial species with varying patterns of pangenome diversity. *Streptococcus pneumoniae* is a nasopharyngeal commensal and pathogen that exchanges genetic material through homologous recombination and has a relatively small core and large accessory genome (~1000 core genes, 5000-7000 accessory genes (Hiller & Sá-Leão, 2018)). *Mycobacterium tuberculosis* is a slow-replicating respiratory pathogen with a low mutation rate and a small accessory genome (~4000 core genes, <1000 accessory genes (Yang *et al*., 2018)). *Escherichia coli* is a genetically diverse enteric bacterium, with an intermediate-size core genome, but extensive accessory genome (~3000 core genes, >100,000 accessory genes (Park *et al*., 2019)). These three species are important, commonly studied pathogens that represent a broad range of pangenome diversity, providing a diverse benchmarking dataset. Genomes for *M. tuberculosis* (N=219), *S. pneumoniae* (N=616) and *E. coli* (N=162) were collated from public repositories (See **Methods**). The same workflows as above were compared based on respective predicted COG frequencies.

Predicted gene frequency histograms for each species and workflow are shown in Figure 5, with counts of COGs with 100% frequency, <100% frequency and total COGs in each pangenome in Table 1. For *M. tuberculosis*, all tools predicted between 3813-3996 COGs at between 90-100% frequency. All tools, except for Roary, predicted a minimal accessory genome, with ggCaller predicting the fewest COGs in the lowest gene frequency bin (66, 0-10% frequency). This is consistent with previous analysis using Panaroo (Tonkin-Hill *et al*., 2020). Notably, Roary predicted the highest number of COGs in the lowest bin (1096, 0-10% frequency) and the highest number of total COGs, likely due to its strict clustering threshold.

**Figure 5:**
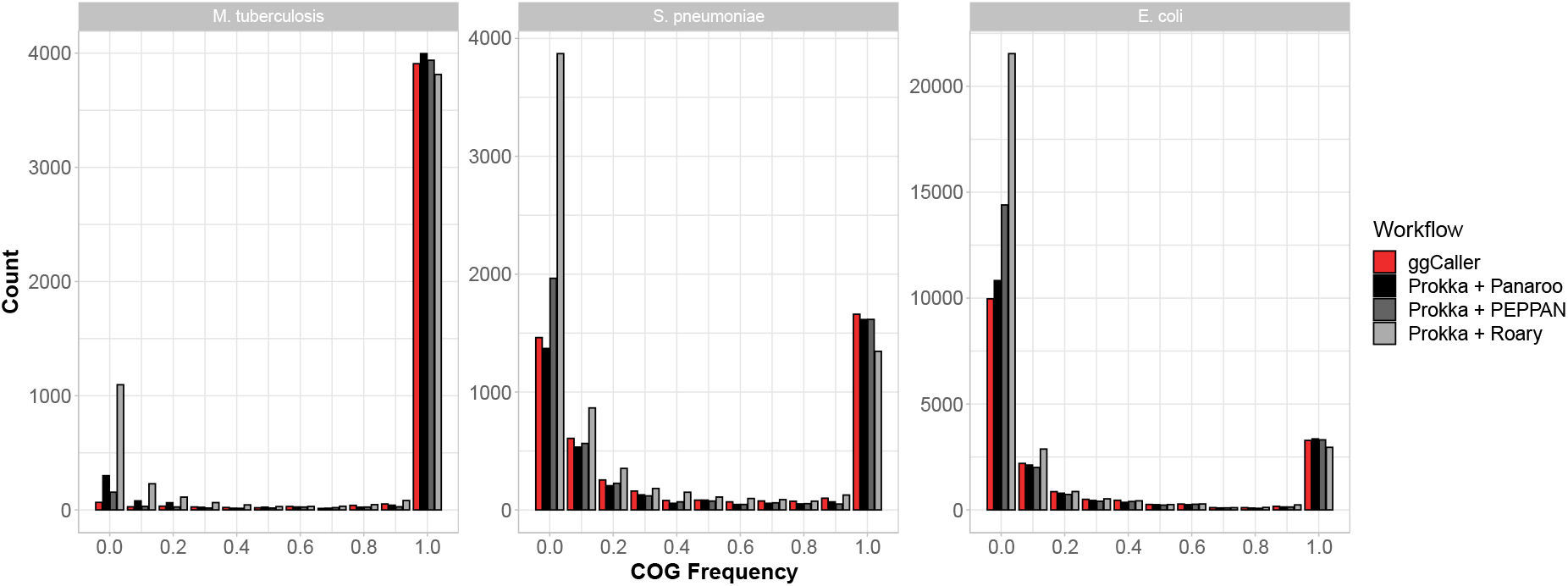
Gene frequency histograms for Mycobacterium tuberculosis, Streptococcus pneumoniae and Escherichia coli across pangenome analysis workflows. ggCaller and Panaroo were run in moderate mode.

**Table 1:**
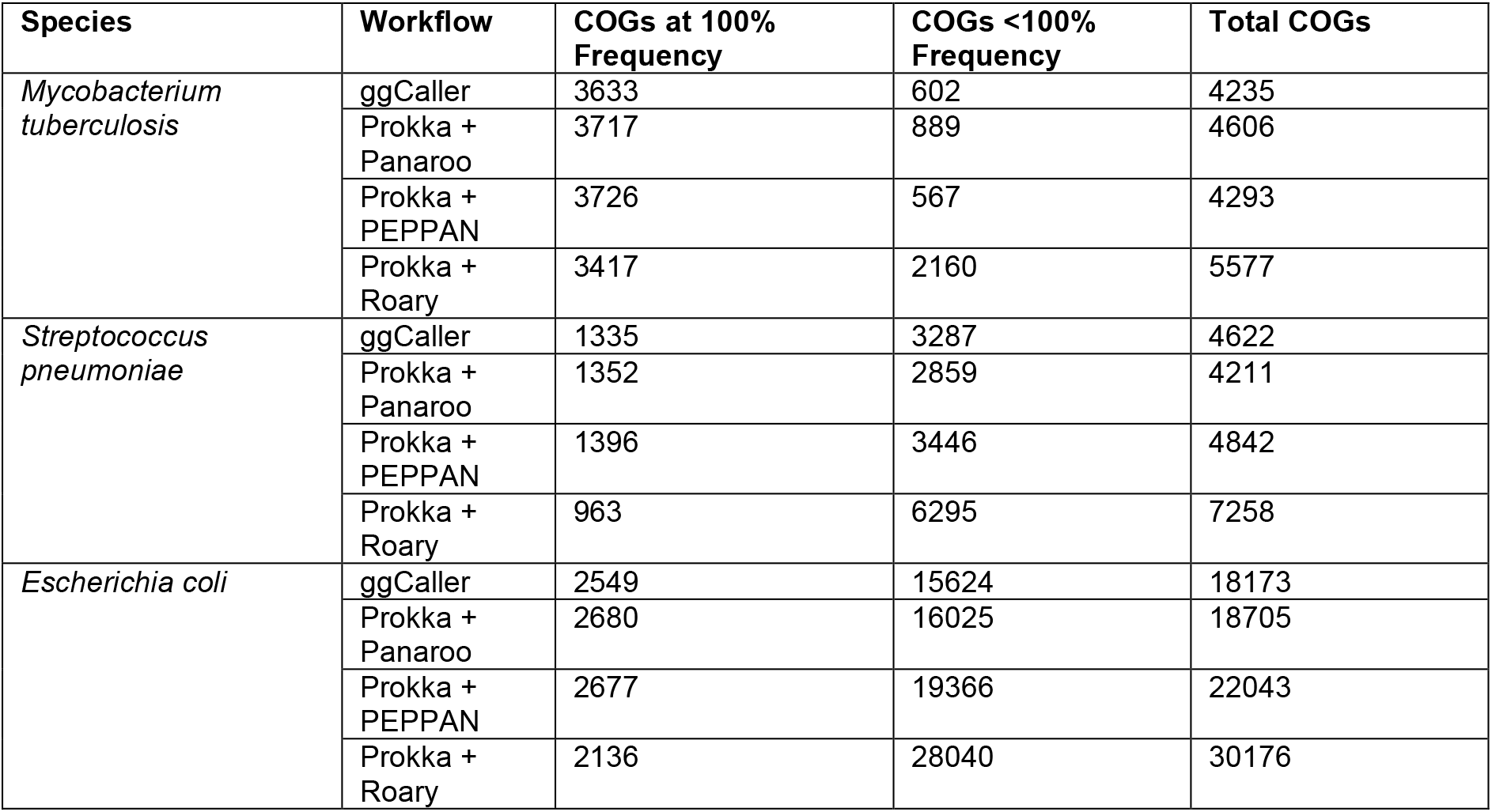
Counts of COGs found at 100% frequency, COGs <100% frequency and total COGs predicted for Mycobacterium tuberculosis, Streptococcus pneumoniae and Escherichia coli across pangenome analysis workflows.

For *S. pneumoniae*, ggCaller, Panaroo and PEPPAN predicted 1660, 1615 and 1616 COGs at between 90-100% frequency respectively, whilst Roary predicted 1345 COGs for the same bin. For the same dataset, Croucher *et al*., (2013a) predicted 1194 COGs at 100% frequency and 5442 total COGs. All tools except for Roary predicted a slightly higher number of COGs found at 100% frequency, and lower total COGs. These differences are likely due to more accurate orthologue clustering compared with the original study, which employed a combination of manual steps and COGsoft (Kristensen *et al*., 2010). More accurate clustering would both reduce the total number of unique COGs reported, and increase the number of high frequency COGs, as seen here for ggCaller, PEPPAN and Panaroo. For low frequency COGs, ggCaller and Panaroo estimated a similar number between 0-10% frequency (1462 and 1370 respectively), with PEPPAN predicting a greater number (1965). Again, Roary estimated the largest number of COGs in all bins between 0-10% frequency (3871) and estimated the largest number of total COGs of all workflows.

For *E. coli*, ggCaller, PEPPAN, Panaroo and Roary estimated a similar number of genes with frequencies in the range 90-100% (3281, 3354, 3308 and 2953 respectively). Estimates of the *E. coli* core genome vary depending on the composition and size of the dataset being analysed, and has previously been reported to be in the range 800-3000 COGs (Kallonen *et al*., 2017; Park *et al*., 2019; Chen *et al*., 2006). Therefore, all predictions were within the expected range for *E. coli*, despite Roary predicting ~300 fewer COGs than the other workflows. ggCaller predictions were consistent with Panaroo and PEPPAN for all frequency compartments, except for COGs found at 0-10% frequency, where PEPPAN predictions were elevated, as seen before with *S. pneumoniae*. This was also consistent with previous simulation results with PEPPAN **(Figure 4B**).

In this analysis of real bacterial populations, ggCaller performed equivalently to existing gold-standard pangenome analysis tools across a broad range of bacterial species, and provides gene frequency predictions in line with previous studies.

### ggCaller annotates structurally complex and repetitive genes more accurately than existing tools

Previous analysis in simulated populations highlighted that increased within-gene divergence can impact estimates of pangenome size and COG annotation **(Figure 3A**). Therefore, genes may be inaccurately annotated or clustered by current approaches due to sequence or structural diversity, for example in antigens under diversifying selection (Croucher *et al*., 2017). To determine the effect of allelic and structural variation within genes on clustering accuracy in ggCaller and alternative tools, we compared examples of structurally diverse COGs from the *S. pneumoniae* dataset used previously. Four structurally diverse proteins were chosen; penicillin binding proteins 1a and 2b (Pbp1a, Pbp2b), Pneumococcal surface protein A (PspA) and Pneumococcal Serine-Rich Repeat Protein (PsrP). Pbp1a and Pbp2b are clinically important due to conferral of beta-lactam resistance and vary structurally through interspecies recombinations, generating mosaic sequences (Croucher *et al*., 2013a). PspA is an important virulence factor under positive selection by the immune system, which has generated wide structural diversity. Finally, the presence of repeats in PsrP (>1000 repeats of SASX motif (Shivshankar *et al*., 2009)) presents a particular challenge for assemblers and gene prediction tools (Croucher *et al*., 2017).

To benchmark the annotation and clustering accuracy of these genes, we compared workflows based on consistency of predicted start and stop coordinates, sequence identity, and the total number of sequences within each COG. As a benchmark, predictions were also compared to the original predicted protein sequences from Croucher *et al*., (2015), where genes were predicted using multiple gene-callers followed by manual inspection to increase accuracy and clustered using COGsoft (Kristensen *et al*., 2010) (referred to as ‘ M anual + COGsoft). Protein sequences from predicted genes were aligned to manually curated reference sequences from Spn23F. Differences between the start and stop positions were compared using the number of amino-acids soft-clipped at either end of the alignment to Spn23F. Soft-clipping is a measure of the ‘over-hang’ of an alignment, with bases in a soft-clip not being aligned to amino acids in the other sequence. Here, a positive value means the query sequence is soft-clipped, a negative value means the correct sequence is soft-clipped, and a value of zero means perfect alignment **(Figure 6A**). Pairwise average amino acid identity (AAI) was also calculated for all sequences within each COG (proportion of matching amino acids over the gapped alignment length (Doolittle, 1981; Raghava & Barton, 2006)). Comparing distributions of AAI across tools provides a measure of clustering accuracy; low peaks indicate start or stop sites are inconsistently predicted between orthologues, whilst high peaks suggest good consistency in gene prediction within a COG. The numbers of sequences within each COG were also considered, to ensure absence of low AAI peaks was not a consequence of COGs containing fewer sequences.

**Figure 6:**
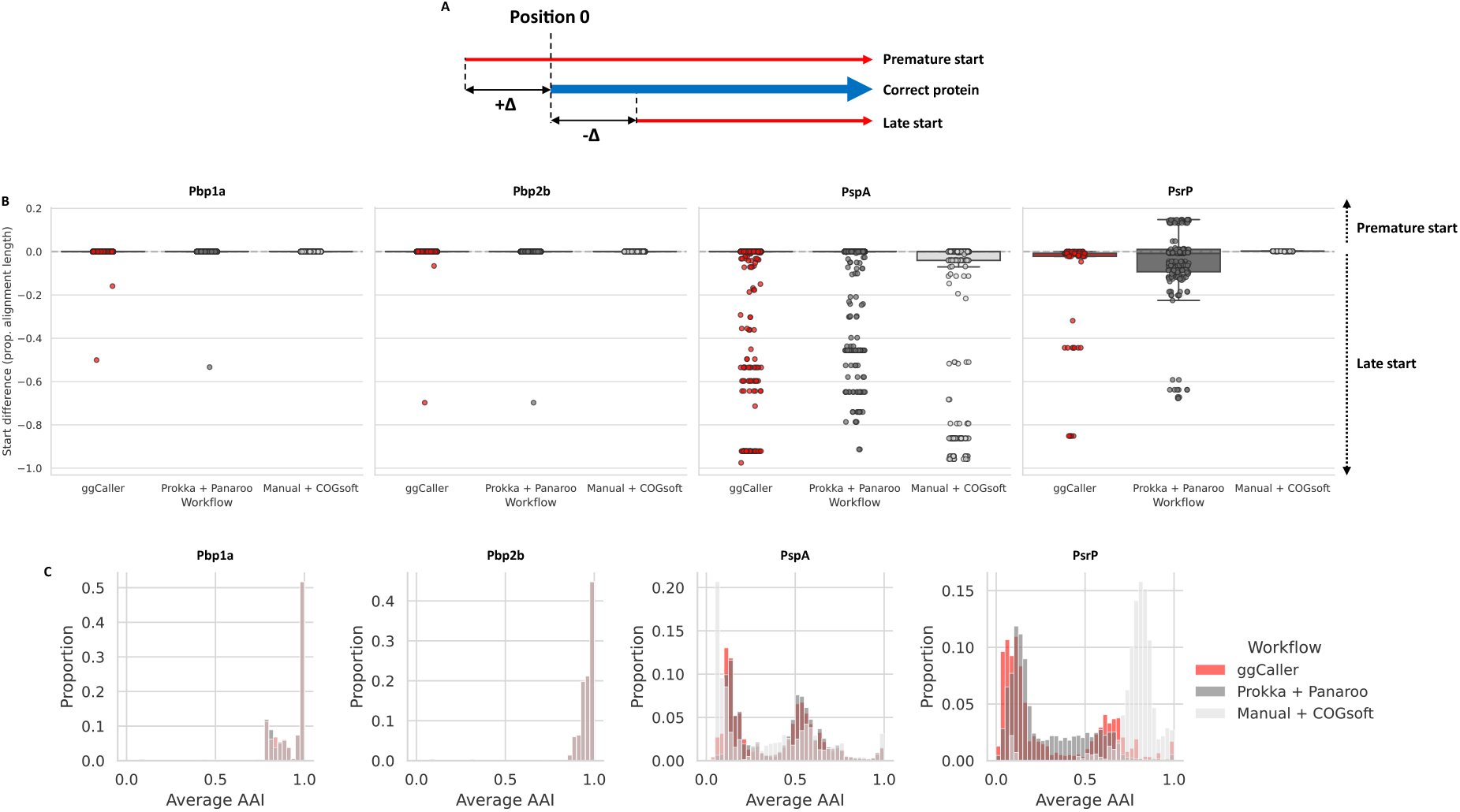
Comparison of within-COG start site soft-clipping and average amino acid identity (AAI). Description of start site soft clipping (**A**). Comparison of start site soft-clipping protein sequences of Pbp1a, pbp2a, PspA and PsrP based on alignment with the manually annotated reference in Spn23F (**B**). Histograms of pairwise average AAI within each COG (**C**).

Comparisons of start site soft-clipping between ggCaller, Prokka + Panaroo and the clustering data from the original study (Manual + COGsoft) are shown in **Figure 6B**, and distributions of pairwise average AAI are shown in **Figure 6C**. For Pbp1a and Pbp2b, almost all predicted proteins matched the start positions within the Spn23F reference for ggCaller and Prokka + Panaroo, and were consistent with Manual + COGsoft. Both workflows also had equivalent distributions of AAI and matched Manual + COGsoft with modal peaks at 1.0, indicating consistent annotation and clustering of orthologues. The number of Pbp1a and Pbp2b orthologues, and the genomes they were found in, also matched Manual + COGsoft (Table 2).

**Table 2:** Number of sequences of Pbp1a, Pbp2b, PspA and PsrP identified and genomes in which they were found for across 616 Massachusetts S. pneumoniae genomes.

| Protein | Workflow/Dataset | No. sequences | No. genomes (% of dataset) |
| --- | --- | --- | --- |
| Pbp1a | ggCaller | 617 | 616 (100%) |
|  | Prokka + Panaroo | 616 | 616 (100%) |
|  | Manual + COGsoft | 616 | 616 (100%) |
| Pbp2b | ggCaller | 617 | 616 (100%) |
|  | Prokka + Panaroo | 616 | 616 (100%) |
|  | Manual + COGsoft | 616 | 616 (100%) |
| PspA | ggCaller | 621 | 548 (89.0%) |
|  | Prokka + Panaroo | 574 | 527 (85.6%) |
|  | Manual + COGsoft | 444 | 420 (68.2%) |
| PsrP | ggCaller | 90 | 84 (13.6%) |
|  | Prokka + Panaroo | 210 | 169 (27.4%) |
|  | Manual + COGsoft | 66 | 66 (10.7%) |

For PspA, distributions of start site variants, as well as AAI distributions, were consistent between ggCaller and Prokka + Panaroo, but were variable within respective COGs. Therefore, ggCaller and Prokka + Panaroo identified similar levels of diversity in PspA. For Manual + COGsoft, there were a greater proportion of truncated proteins present than both the other workflows, and although its respective AAI distribution was largely consistent with ggCaller and Prokka + Panaroo, fewer PspA orthologues were identified. PspA is a core gene in *S. pneumoniae* (Croucher *et al*., 2017), and therefore should be identifiable in all isolates. ggCaller had greater recall than Prokka + Panaroo and Manual + COGsoft, identifying PspA in 21 and 128 more isolates respectively.

For PsrP, ggCaller gene annotations had lower start site soft-clipping compared to Prokka + Panaroo, and were consistent with Manual + COGsoft. Both ggCaller and Prokka + Panaroo PsrP annotations had broad multi-modal distributions of AAI, whilst Manual + COGsoft had a single modal peak at ~80% AAI. However, PsrP stop site predictions with Manual + COGsoft were almost all truncated, whilst ggCaller predictions matched Spn23F (**Supplementary Figure 4**). This discrepancy explains the higher overall AAI for Manual + COGsoft: fewer repeat units were included in PsrP sequences, leading to more closely matching sequences, whilst ggCaller correctly predicted more gene end coordinates. Both ggCaller and Prokka + Panaroo had a modal peak at ~25% AAI. However, ggCaller identified proteins with a greater proportion spanning 50-100% AAI than the other methods. From the raw FASTA files, only 15/210 PsrP sequences contained an SASX motif for Prokka + Panaroo, compared to 39/90 and 66/66 for ggCaller and COGsoft respectively. The DAE motif, present in incorrect CDSs originating on the reverse strand to *psrP*, was found in 148/210 and 46/90 sequences for Prokka + Panaroo and ggCaller respectively, whilst it was not found in any for Manual + COGsoft. Whilst these results highlight issues with gene prediction for both automated processes, ggCaller still identified more correct PsrP sequences. ggCaller was also consistent with Manual + COGsoft in terms of COG size, which identified PsrP in 84 and 66 genomes respectively, versus 169 in Prokka + Panaroo. Due to the poor alignment of start sites and modal peaks at low AAI, Prokka + Panaroo likely inflated the size of the PsrP COG due to mis-clustering of incorrectly extended and truncated sequences. Overall, ggCaller outperformed a workflow containing current gene annotation and pangenome analysis tools when clustering structurally diverse proteins.

### ggCaller improves functional interpretation in pangenome-wide association studies

ggCaller supports querying of sequences of arbitrary length within an annotated DBG, enabling reference-free functional interpretation of sequence elements. This is useful when analysing significant hits from a pangenome-wide association study (PGWAS), where current approaches annotate results by mapping to only one or a few references (Lees *et al*., 2018). To demonstrate the utility of this approach, we performed PGWAS to identify sequences significantly associated with tetracycline and macrolide resistance in *S. pneumoniae*. Tetracycline resistance is caused by presence of *tetM* in *S. pneumoniae*, which is associated with conjugative transposon Tn*916* (Croucher *et al*., 2009). Macrolide resistance can be caused by presence of *mef/mel*, which are found on a variety of gene cassettes that integrate at multiple sites around the genome (D’Aeth *et al*., 2021). These genes are not present in all *S. pneumoniae* isolates (Croucher *et al*., 2013a), therefore annotation accuracy of significant hits will depend on presence of the gene in a chosen reference. Consequently, correct interpretation of macrolide resistance PGWAS in this species has proved challenging using previous approaches (Lees *et al*., 2016). To highlight issues with using a single reference for annotation, unitigs associated with either tetracycline or macrolide (represented here by erythromycin) resistance were identified in 616 *S. pneumoniae* genomes with comprehensive minimum inhibitory concentration (MIC) data (Croucher *et al*., 2015) using pyseer (Lees *et al*., 2018). A core genome phylogeny was generated using ggCaller for each antibiotic dataset and used for pyseer population-structure correction (**Supplementary Figure 5**). This phylogeny highlighted a correlation between gene presence, AMR phenotype and population structure, as seen in Croucher *et al*., (2013a). Annotations of significant unitigs were compared between ggCaller and the built-in pyseer annotation function using only Spn23F as a reference.

This PGWAS identified a total of 1517 and 703 significant unitig hits for tetracycline and erythromycin resistance respectively. Mapping these hits to a single reference (Figure 7A) showed a strong signal at Tn*916* (peak 3) for tetracycline resistance, which contains *tetM*. In contrast, two weaker signals were present at loci associated with erythromycin resistance (peaks 1 and 2). Based on Spn23F annotation, peak 1 aligns to a locus containing the glycosyltransferase, *capD* (locus tag: Spn23F01130), whilst peak 2 aligns to a locus containing the DNA-3’-methyladenine glycosylase I, *tag* (locus tag: Spn23F01760). Both loci have been identified as insertion sites for *Tn1207.1-type* elements that can harbour *mef/mel* genes (D’Aeth *et al*., 2021). As Spn23F does not contain *mef/mel*, these peaks are false positive hits resulting from linkage disequilibrium between orthologues of *capD* and *tag*, and loci associated with erythromycin resistance. In contrast, mapping significant unitigs to DBGs annotated by ggCaller correctly and directly identified the causal genes. Genes annotated as *tetM* had the greatest coverage of significant unitigs associated with tetracycline **(Figure 7B**), whilst genes annotated as *mefE* were top for erythromycin, with *mel* having the 6^th^ highest coverage **(Figure 7C**). Therefore, ggCaller provides a useful extension to PGWAS to avoid incorrect or difficult manual functional inference of hits when restricted by arbitrary reference genome choice.

**Figure 7:**
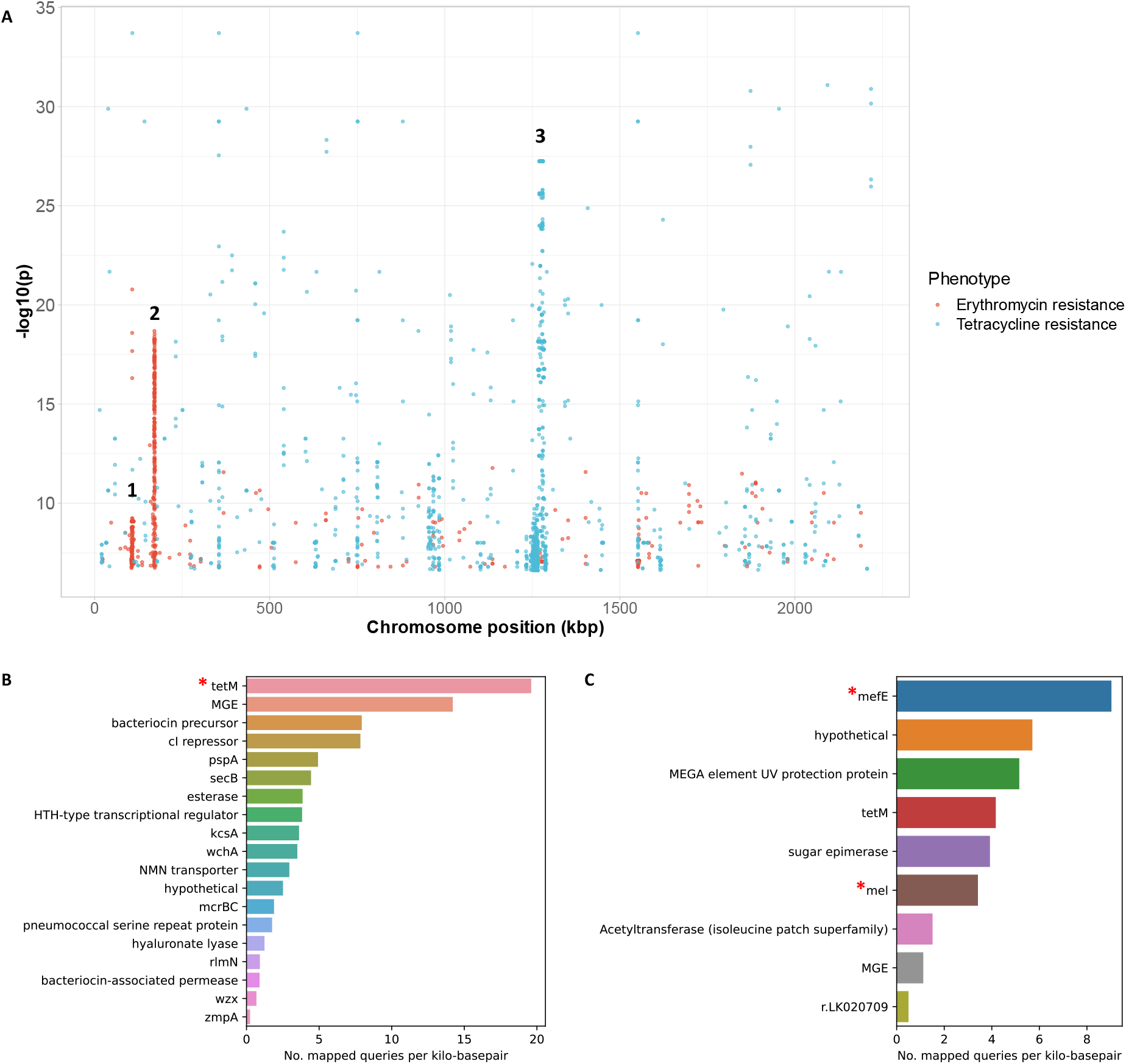
Pangenome-wide association study of tetracycline and erythromycin resistance. Manhattan plot of unitigs mapped to Spn23F. Locus tags and gene names for features in peaks; 1: Spn23F01130 (*capD*), Spn23F01140 (*epsC*), 2: Spn23F01760 (*ruvA*), Spn23F01770 (*tag*), Spn23F01780 (putative protease), 3: Tn*916* (**A**). Gene coverage by significant unitigs associated with tetracycline (**B**) and erythromycin (**C**) resistance identified by pyseer and matched by ggCaller, with known causative genes marked by red asterisks. Number of mapped queries per kilo-basepair was calculated by binning genes matched to queries by ggCaller with the same annotation, and then taking the ratio of the number of queries mapped to the total sequence length of the bin. Core genome phylogenies with resistance and causal gene annotations generated by ggCaller are available in **Supplementary Figure 5**.

### ggCaller is faster than per-genome annotation and clustering

Existing pangenome analysis workflows rely on iterative and usually redundant annotation of genes within independent genomes. In contrast, ggCaller simultaneously annotates genes across a population within a DBG. Therefore, ggCaller computational performance is expected to scale with DBG complexity (given by number of nodes and edges), rather than linearly with the number of samples. To understand the effect DBG complexity has on computational performance, we benchmarked ggCaller against Prokka + Panaroo using two *S. pneumoniae* datasets with different levels of pangenome diversity. Genomes from the worldwide Global Pneumococcal Sequencing project (Gladstone *et al*., 2019) and the statewide Massachusetts dataset from Croucher *et al*., (2015) were used to represent variable levels of diversity within a single pathogen. The number of nodes and edges in DBGs for the global dataset were greater than the Massachusetts dataset for the same number of isolate genomes (**Supplementary Figure 6**), highlighting greater diversity in the global dataset. To ensure consistent annotation across workflows, the same CDS annotation database from the Massachusetts study was provided to both Prokka and ggCaller.

ggCaller runtime was greatly reduced compared to Prokka + Panaroo for both the global and Massachusetts datasets **(Figure 8A**), even as dataset size increased. For example, at 100 genomes, ggCaller had 30.3x and 61.9x speed-up vs Prokka + Panaroo for the global and Massachusetts dataset respectively, decreasing to 7.5x and 47x at 500 genomes, and 3.5x at 1000 genomes for the global dataset only (**Supplementary Figure 7**). For Prokka + Panaroo, the bulk of processing time was spent during Prokka gene annotation, consistent with other workflows using Prokka with Roary and PEPPAN (**Supplementary Figure 8**). This process relies on individual annotation of genes in each genome through BLAST and HMM search resulting in repeated computation, rather than sharing annotations across orthologues in ggCaller.

**Figure 8:**
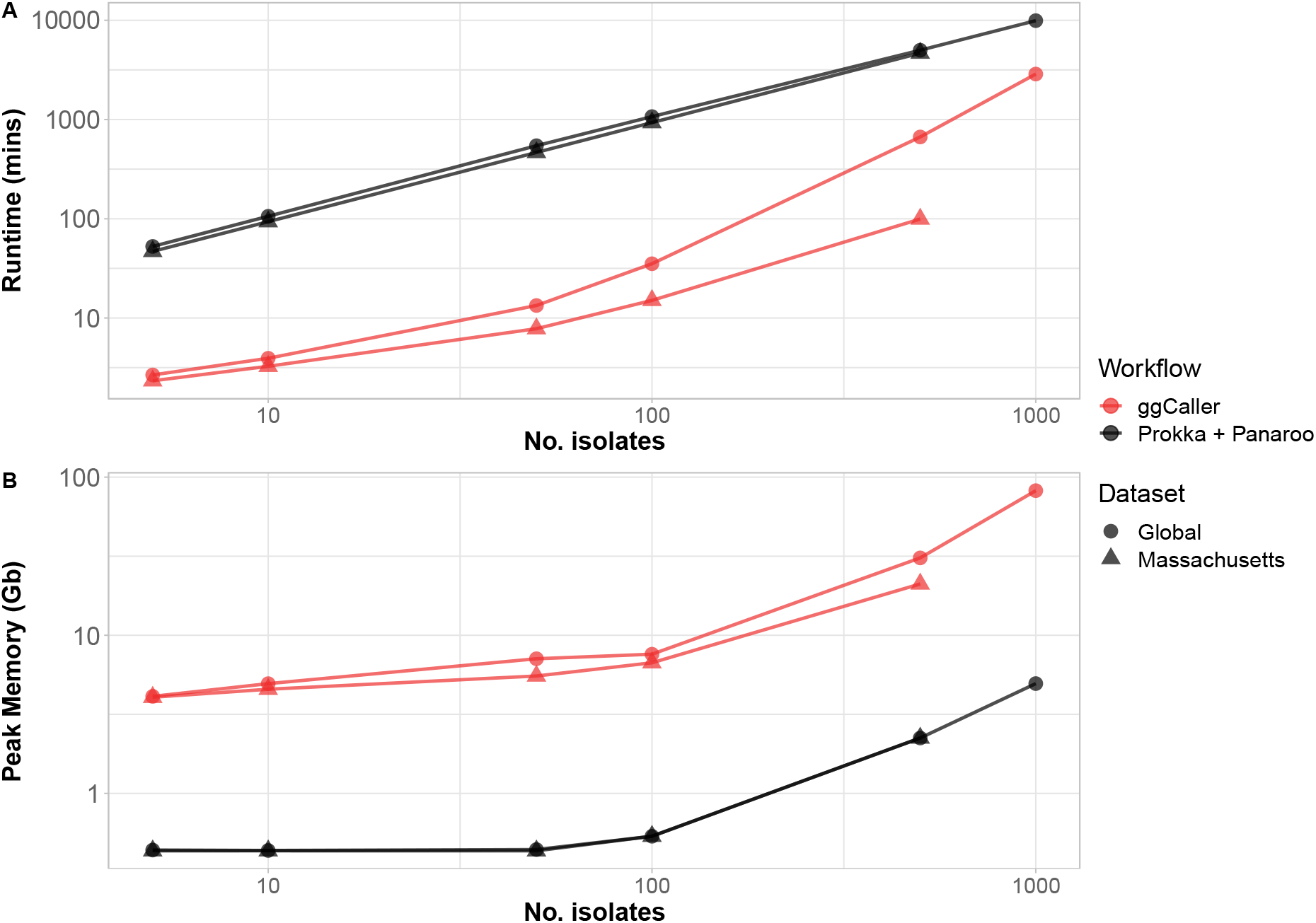
Computational benchmarking of ggCaller against Prokka + Panaroo. Tools were run using 16 threads, comparing runtime (**A**) and peak memory (**B**), with an increasing number of randomly sampled genomes.

Conversely, ggCaller memory use was higher than Prokka + Panaroo **(Figure 8B**). This is due to ggCaller storing the DBG and gene-calls in memory for fast access, leading to increasing memory usage as the population complexity increases. ggCaller memory usage was lower for the Massachusetts than the global dataset for >10 genomes. Furthermore, both runtime and memory usage scaled with the number of nodes and edges within the DBG (**Supplementary Figure 6**), indicating graph complexity, and by extension pangenome diversity, impacts performance. Overall, ggCaller shows greatly reduced runtime versus an existing pangenome analysis workflow, with its performance scaling with diversity.

## Discussion

In recent years, there has been increased focus on improving the accuracy, functionality and sensitivity of bacterial gene annotation, as well as the overall usability of software tools. Prokka (Seemann, 2014), DFAST (Tanizawa, Fujisawa & Nakamura, 2018) and Bakta (Schwengers *et al*., 2021) were all developed over the last decade as stand-alone tools that combine gene prediction and functional annotation. However, innovation in the underlying algorithms for gene prediction has stalled; all of the above tools rely on Prodigal for gene prediction (Hyatt *et al*., 2010). Moreover, bacterial genomes are now no longer analysed in isolation; datasets of hundreds or thousands of sequences are routinely generated and analysed at once (Land *et al*., 2015). Existing pangenome analysis tools already use information provided from simultaneous analysis of many genomes to improve accuracy (Tonkin-Hill *et al*., 2020; Zhou, Charlesworth & Achtman, 2020). However, the upstream process of bacterial gene annotation is still conducted on individual genomes. Therefore, there is huge redundancy and potential for mis-annotation when annotating the same gene across multiple genomes. These issues lead to longer runtimes and inaccurate clustering, ultimately impacting inferences made on population structure and gene distributions (Dimonaco *et al*., 2022; Tonkin-Hill *et al*., 2020; Zhou, Charlesworth & Achtman, 2020).

We developed ggCaller to leverage population-frequency information to improve the accuracy and speed of gene identification, annotation and pangenome analysis. ggCaller predicts and annotates genes within a pangenome de Bruijn Graph (DBG) built from hundreds of individual genomes. Sequence sharing, encoded as node frequencies by the DBG, enables several innovations in ggCaller over existing tools: **i**) contig breaks can be traversed using identical paths present in other assemblies, **ii**) ORF start site frequencies are used to consistently predict start codons, **iii**) ORF scores generated by BALROG temporal convolutional networks (Sommer & Salzberg, 2021) are shared across COGs during ORF filtering, **iv**) genes are functionally annotated within COGs, and **v**) an updated version of Panaroo is implemented for iterative gene clustering, paralogue identification, removal of erroneous CDSs and re-identification of genes missed on the first pass.

ggCaller outperformed existing state-of-the-art tools when applied to a diverse set of simulated and real bacterial datasets. Gene predictions were more consistent in terms of start and stop codon identification, and within-COG sequence identity, leading to more accurate clustering and gene frequency distributions. ggCaller was also less sensitive to highly fragmented assemblies than existing tools, enabling greater recall of full-length genes. In terms of computational performance, ggCaller had a largely reduced runtime against a workflow of Prokka and Panaroo by removing redundancy in scoring and annotation.

ggCaller is a useful addition to pangenome-wide association studies (PGWAS), enabling reference-agnostic functional annotation when used alongside tools such as pyseer (Lees *et al*., 2018) or DBGWAS (Jaillard *et al*., 2018). ggCaller has a streamlined workflow for DBG annotation, core-genome phylogeny generation and significant hit annotation. When applied to *S. pneumoniae* PGWAS of two AMR phenotypes, ggCaller provided a simple, accurate functional interpretation of significant hits. In contrast, using a single reference required expert knowledge of the species’ genome biology and relevant literature, and highlighted that functional interpretation of significant hits can be greatly affected by choice of reference sequence. By extension, ggCaller can be used by any study linking sequence to phenotype, such as in pangenome-wide epistasis analysis (Pensar *et al*., 2019), or in development of models for phenotype prediction from genomic data (Lees *et al*., 2020).

A technical limitation of the current version of ggCaller is its memory usage, as the DBG and all gene-calls across the population are stored for fast access. However, both runtime and memory usage varied depending on choice of dataset. Pangenome diversity is a key factor in ggCaller scalability, as including more variation will increase DBG complexity. A less diverse dataset (e.g., a single sequence type or clonal complex) will see the greatest improvement in runtime with ggCaller over current state-of-the-art workflows, alongside less extreme memory usage. More diverse datasets (e.g., global collection of sequence types) will still likely see a speed-up using ggCaller, albeit the effect will be reduced. This scaling with graph complexity places ggCaller in unique position amongst pangenomic analysis workflows, meaning it is well-suited for analysis of more-closely related isolates, such as in regional surveillance. Further work will aim improve scalability, particularly with memory usage, which can be addressed using memory mapping to leverage low-latency storage media.

Additionally, ggCaller cannot yet be run iteratively, requiring the full complement of genomes to be supplied at the start of analysis. This ‘online’ functionality is a desirable feature for epidemiological tools, as new genomes will inevitably be added to datasets. Moreover, an alternative function of ggCaller is gene prediction in unassembled datasets, as Bifrost DBGs can be built from reads (Holley & Melsted, 2020). However, graphs from read data are complex, and contain paths that do not represent real sequences, so this was not tested here. Finally, ggCaller is limited to identification of bacterial coding sequences, meaning annotation of non-bacterial genes and non-coding RNA is not currently supported.

ggCaller is a novel bacterial gene annotation and pangenome analysis tool which outperforms existing state-of-the-art tools in terms of both speed and accuracy, achieved through its use of pangenome de Bruijn graphs. ggCaller also enables reference-agnostic functional inference, making it an important extension to pangenome-wide association studies. Graph-based analysis has the potential to become the new convention in bacterial genomics, bringing with it benefits of reduced redundancy, increased consistency and improved accuracy over linear-genome based methods. Enabling graph-based annotation and pangenome analysis is an important step in this transition.

## Methods

### Code availability

ggCaller source code is available at https://github.com/samhorsfield96/ggCaller under the open-source MIT license. All analysis scripts, instructions on how to use them, and summary data are available at https://github.com/samhorsfield96/ggCaller_manuscript. ggCaller v1.3.3 was used for all analysis and is available as a release on Github (https://github.com/samhorsfield96/ggCaller/releases/tag/v1.3.3).

### Graph indexing

The de Bruijn graph (DBG) used by ggCaller is generated by Bifrost (Holley & Melsted, 2020). Bifrost was chosen as it is highly scalable to hundreds of thousands of bacterial genomes and has an intuitive C++ API. To build a DBG, Bifrost first generates an index of all k-mers in the population using a blocked Bloom filter. *k* = 31 has been shown to provide a good balance of efficiency and sensitivity of detection of shared/divergent sequences in bacterial genomes (Holley & Melsted, 2020), and so is used as the default in ggCaller. Starting with a single k-mer, Bifrost queries the presence of its suffix (length= *k* – 1) appended with each of A, C, G and T within the blocked Bloom filter (Figure 1, **step 1**). Connections of length *k-1* between k-mer suffixes and prefixes are treated as edges. Each k-mer is assigned colours, which describe in which genomes they are present in. Unitigs, also known as nodes, are generated by merging unbranching k-mer paths into a single DNA sequence. Branching paths, described by edges, represent variation within a population. A genome can be regenerated from a DBG by traversing a path made up of nodes assigned with a specific colour.

Once the DBG is built, ggCaller iterates over the DNA sequence in each node and identifies stop codons in each reading frame, storing this information in a six-bit bitset, where each bit represents a reading frame (three forward and three reverse) (Figure 1, **step 2**). A bit is set to ‘1’ if at least one stop codon is present in that reading frame, and ‘0’ otherwise. ggCaller stores the node colours in a separate bitset; as the colours of the constituent k-mers within a node may not all be identical due to contig breaks (Schulz, Wittler & Stoye, 2022), ggCaller takes the intersection of the colours of the start and end k-mers for each node. ggCaller additionally determines the frequency of stop codons for each reading frame within the graph by counting the number of set bits. For accurate start site identification, ggCaller also identifies the first base of all start codons, and *k* – 1 downstream bases (*k* chosen as this is the shortest possible node length). This sequence is then translated into its respective amino acid sequence and further split into l-mers (*l* = *m* / 3, where *m* = Bifrost minimiser size, chosen to ensure sufficient coverage of each start site). The coverage of each l-mer is calculated within the population and stored in a robin-hood unordered map to be used using graph traversal. Using amino-acid l-mers reduces the impact of small mutations on the calculation of start site frequency, which would otherwise make start sites of non-identical orthologues appear less frequent.

As Bifrost does not consider sequence contiguity from input genomes when inferring edges, traversal of a Bifrost DBG can result in generation of sequences that are not present in the original genomes (Břinda, Baym & Kucherov, 2021). To prevent this, candidate paths are queried against input genomes to ensure they are present as contiguous sequence in the input. An FM-index is built from each genome in node space (using node identifiers rather than DNA sequence) using kseq (https://github.com/lh3/seqtk) and SDSL v3 (Gog *et al*., 2014), to which paths can be compared during traversal. Building an FM-index using node identifiers rather than DNA sequence greatly reduces the size of the index, reducing the memory and time required to load and search the index.

### Stop codon search using iterative graph traversal

ggCaller uses an iterative depth-first search (DFS) algorithm to pair stop-codons in the same reading frame (Figure 1, **step 3a**). To reduce the search space, ggCaller traverses the DBG colour-by-colour (i.e. sample-by-sample). Starting at a source node containing at least one stop codon, ggCaller iteratively visits all neighbour nodes with the same colour, with each neighbour added to the top of a stack. Each iteration of the algorithm removes the top node from the stack and visits its neighbours with the same colour as the source node. As each neighbour is visited, the complete path in node space is queried against an FM-index of the given colour to ensure the path is also contiguous in the underlying genome (Figure 1, **step 3b**). This process is repeated until all stop codons in the source node have been paired with a downstream stop codon in the same reading frame. This is now designated as a complete ‘stop-to-stop’ path. This process is repeated for each node containing at least one stop codon, and for each colour in the DBG.

### ORF identification within stop-to-stop paths

For each complete ‘stop-to-stop’ path, ggCaller identifies all possible start codons in the path and pairs them with their respective downstream stop codons to generate a list of candidate ORFs (Figure 1, **step 4**). Starting at the start site for the longest possible ORF, start positions for consecutively shorter ORFs are compared to the current ‘high-scorer’, with the score dictated by: **i**) the translation initiation site score given by BALROG, **ii**) the median population coverage for all amino-acid l-mers in the start site, and **iii**) the number of times the start site has been chosen as the high-scorer in orthologous ORFs. For a start position of a shorter ORF to be chosen as a high-scorer, it must ‘win’ in 2/3 of the above criteria against the previous high-scorer.

To avoid checking start positions of short ORFs that are likely to be incorrect, their start sites are penalised using a probabilistic model. As genes are characterised by long stretches of DNA without two stop codons in the same reading frame, the average frequency of stop codons within genes will be lower than the population average. Therefore, short ORFs can be penalised based on the probability of observing the longer of two ORFs if stop codons occurred at a constant rate (**Equation 1**). This is given by the population stop-codon frequency (p) and the difference in length between the long (*l*0) and short (*l*1) ORF. This gives the probability of the longer ORF occurring by chance if stop codons occurred randomly at the population average frequency (*q*). *q* is only calculated between the current high-scorer and next start site in the iteration. If *q* falls below a threshold, meaning the probability of observing the current high scorer is unlikely by chance, all subsequent shorter ORFs are ignored. A default of *q* < 0.2 (i.e. <20% probability of longer ORF occurring by chance) was chosen, as this provides a good trade-off between sensitivity and precision.

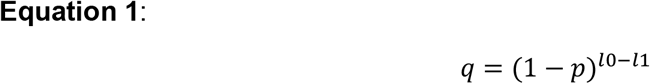

### ORF clustering and gene scoring

At this stage, all ORFs have been identified across all colours. However, many of these sequences are similar, so to remove redundancy from downstream scoring and pangenome steps, ggCaller first clusters ORFs using an algorithm with a similar approach to Linclust (Steinegger & Söding, 2018), but adapted for DBG sequences (Figure 1, **step 5a**). Firstly, ggCaller identifies all sets of ORFs which share at least one node as candidate clusters, placing them in a ‘node group’, and identifies the largest ORF for each node group as a centre sequence. ggCaller then translates every ORF and aligns them with the centre sequence of the node group(s) it belongs to using Edlib (Šošić & Šikić, 2017). If the comparison surpasses a given identity and length cut-off, the ORF is clustered with the centre sequence. A default of 98% for identity and length cut-off was chosen, as these are the initial clustering parameters used in Panaroo for identifying orthologues. For ORFs that do not cluster with a centre sequence, the process is repeated until each ORF is either paired with a centre sequence, or is a centre sequence itself.

Centre sequences are then scored based on how likely they are to be a real coding sequence by using BALROG (Figure 1, **step 5b**). BALROG implements a temporal convolutional network which generates an average score per amino-acid residue, which is incorporated with start codon sequence, the length and TIS score to generate an individual score for each ORF in the cluster. ORFs with a score below a pre-defined cut-off (default = 100, as used in BALROG) are removed from the set. Those surpassing the score are regarded as candidate protein coding sequences (CDSs).

### CDS filtering to coding sequence identification

In the final prediction step, overlapping CDSs must be filtered to pick the most likely final gene calls. This is achieved by finding the highest-scoring tiling path through regions of the DBG with overlapping CDSs. Overlaps between CDSs are first calculated for each colour using a sparse matrix generated using Eigen v3 (Guennebaud, Jacob *et al*., 2010). Each row in the sparse matrix is an CDS ID and each column is a node in the DBG, with ‘1’ in each cell where the node is contained within the ORF, and ‘0’ otherwise. This matrix is multiplied by the transpose of itself, resulting in a sparse matrix with CDS IDs on both rows and columns, and non-zero values off the diagonal indicating ORFs that share at least one node and therefore have a potential overlap. ggCaller then iterates over these potential overlaps, and identifies whether each pair of CDSs genuinely overlap, or whether they simply share sequence, such as a repeated motif. Overlaps are calculated using respective DBG coordinates of each CDS. If two CDSs do overlap, then the number of overlapping base-pairs is calculated, as well as the direction of the overlap.

For each set of overlaps, ggCaller then generates a subgraph of these CDSs as a directed graph using the boost graph library (Siek, Lee & Lumsdaine, 2002), where each node is an CDS, and each edge is an overlap (Figure 1, **step 6**). Each node is labelled with the respective score for that CDS, and each edge with the overlap penalty calculated from the size of the overlap and the penalty scores provided by BALROG. The graph is then separated into connected components, and each component is traversed using the Bellman-Ford algorithm (Bellman, 1958) to determine the highest scoring path. The Bellman-Ford algorithm requires two small modifications to the graph: all scores are multiplied by −1, and all cycles in the graph are pruned via removal of back edges (Siek, Lee & Lumsdaine, 2002). A transitive closure is then generated, which connects all upstream nodes to all downstream nodes that have a viable path between them. This is to enable skipping of CDSs mid-way within a path if a higher scoring path is present. CDSs present in the highest-scoring path are returned as ‘true’ gene-calls.

### Pangenome analysis using DBG-based Panaroo

Panaroo builds COG graphs and applies filtering criteria to prune false positive gene calls, and gene refinding to mitigate false negative gene calls. We integrated these steps into ggCaller for the same advantages, and to simultaneously and efficiently produce an annotated pangenome matrix. The original version of Panaroo (Tonkin-Hill *et al*., 2020) operates on linear genome annotations; a COG graph is then generated by CD-hit iteratively clustering genes, and adjacency information between genes drawn from input assemblies. As it would be redundant to regenerate this information from the above annotations, we implemented a modified version that works directly with the ggCaller annotated graph (Figure 1, **step 7**). Genes which do not have existing neighbours from the overlap step are connected to a neighbour via a DFS of the DBG for each colour. All connected genes are used to generate a COG graph, where genes are added to the same nodes if they belong to the same clusters calculated previously. We implemented the same three stringency settings as Panaroo for removing spurious COGs. Spurious COGs are identified as those at the end of contigs (i.e. have 0/1 edges), or are in a separate component to the majority of COGs. Each stringency mode removes spurious COGs below a certain population frequency (strict: <5%, moderate: <1%, sensitive: none removed).

We also included several alterations to the original Panaroo algorithm. After the initial collapse of gene families, each gene cluster is functionally annotated (i.e. predicted gene name and function through similarity search) based on alignment of each cluster’s centre sequence to a pre-defined annotation database using DIAMOND (Buchfink, Xie & Huson, 2014) and HMMER3 (Eddy, 2009). This annotation is then shared across all members of the cluster. This cluster-by-cluster annotation approach scales more efficiently than existing linear genome annotation tools when analysing many genomes at once. We provide two default annotation databases; Uniprot (Bateman *et al*., 2021) (queried ‘Bacteria’ and ‘Virus’, only reviewed sequences included) for DIAMOND and HMM profiles from Prokka (Seemann, 2014) for HMMER3. Custom databases can also be provided (amino acid FASTA for DIAMOND, HMM-profiles generated by ‘hmmbuild’ for HMMER3). Gene-refinding, which is implemented in Panaroo to search for missing gene-calls within linear genomes, has been altered to work within the Bifrost DBG. We also simplified within-cluster alignment, replacing the choice of aligners with only MAFFT. We implemented two options for running MAFFT: **i**) default mode, where all sequences are aligned together, and **ii**) reference mode, where centre sequences are aligned together, and then the remainder of sequences are aligned in reference-guided mode. The former is faster when only a few sequences are present in each cluster, whereas the latter can be used to speed-up alignment of large clusters.

### Querying of sequences within an annotated DBG

ggCaller supports querying new sequences against an annotated DBG, which is useful during functional inference of significant hits in pangenome wide association studies. Multiple query sequences can be supplied, which are broken into k-mers and queried against the graph using the Bifrost ‘query’ function. This identifies all node sequences that are found in the query sequence. As all CDSs identified by ggCaller are indexed by node, any that share at least one node with the query sequences are returned. Returned CDSs are output in FASTA format, detailing the query or queries with which they overlap, and the genome(s) in which they are found.

### Output files and plots

By default, ggCaller generates FASTA files of all CDSs in nucleotide and amino-acid format. ggCaller also generates the same files as Panaroo (except for gene_data.csv, a summary of CDSs and their annotation, all of which is included in ggCaller FASTA files), including a gene-presence absence matrix, annotated graph in GML format, and per-cluster alignments in FASTA format. If annotation is specified, ggCaller can also generate GFF files for assemblies included in the DBG. We also implemented pairSNP-cpp (https://github.com/gtonkinhill/pairsnp-cpp) and Rapidnj (Simonsen, Mailund & Pedersen, 2008) to rapidly generate SNP distance matrices, and neighbour-joining Newick trees of all input genomes. ggCaller also generates within-cluster alignments and concatenated core-genome alignments for phylogenetic analysis. Additionally, ggCaller uses SNP-sites (Page *et al*., 2016) to identify single nucleotide polymorphisms in alignments in VCF format. ggCaller also generates summary graphs, including a rarefaction curve for estimating pangenome ‘openness’ (Tettelin *et al*., 2005), and gene frequency distributions (Baumdicker, Hess & Pfaffelhuber, 2010), either in terms of cluster size or population proportion.

### Computational optimisations

ggCaller is written in C++ and python, and is parallelizable through use of OpenMP (Dagum & Menon, 1998) and python’s multiprocessing module. ggCaller relies on robin-hood hash maps for in-memory storage of ORF graph coordinates, which are more time and memory efficient than standard C++ unordered maps (https://github.com/martinus/robin-hood-hashing). ggCaller reimplements the BALROG model of translation initiation site and gene structure (Sommer & Salzberg, 2021). The temporal convolutional network structure and weights are rewritten using torch to enable direct C++ integration (Collobert, Bengio & Mariéthoz, 2002). These models were initially trained using pytorch and compatible only with a python API (Paszke *et al*., 2019), with torch reimplementation improving model querying speed and enabling parrallelisation using shared memory. ORF scores generated by BALROG are stored in a lock-free dictionary which can be accessed by independent threads to avoid rescoring the same sequence. ggCaller builds an FM-index using node identifiers instead of DNA sequence to identify incorrect paths generated during DBG traversal. This significantly reduces time and memory requirements during FM-index querying.

### Bacterial datasets used for benchmarking

Seven simulated populations of 100 genomes were generated using the Infinitely Many Genes simulation model (Baumdicker, Hess & Pfaffelhuber, 2010), with the *Streptococcus pneumoniae* ATCC 700669 serotype 23F (termed ‘spn23F’, Genbank accession: FM211187.1) (Croucher *et al*., 2009) reference genome as the root. This process is available as a custom script, which was used previously in the validation of Panaroo (simulate_full_pangenome.py). Parameters of each simulation are detailed in **Supplementary Table 1**. For the contaminated simulation, random 10 kb fragments of the *Staphylococcus epidermidis* ASM764v1 chromosome (Genbank accession: AE015929.1), which is a common contaminant, were inserted into each assembly. For the fragmented simulation, assemblies were sheared based on real contig fragment lengths from assemblies in Croucher *et al*., (2015). For each simulation, FASTA files containing simulation assemblies and ground-truth CDS annotations were generated. Illumina paired-end reads were then simulated from all assemblies using ART v2.5.8 (Huang *et al*., 2012), and assembled using SPADES v3.15.3 (Bankevich *et al*., 2012).

*Streptococcus pneumoniae* genomes (N=616) were gathered from Croucher *et al*., (2015). A representative subset of genomes from a dataset of *Escherichia coli* (N=162) were gathered from an analysis in Lees *et al*., (2019), originally from Kallonen *et al*., (2017). *Mycobacterium tuberculosis* genomes (N=219) were also gathered from Lees *et al*., originally from Cohen *et al*., (2015).

### Linear-genome gene annotation

For linear-genome based pangenome analysis, genes were called using Prokka v1.14.6 (Seemann, 2014) or GeneMarkS-2 v1.24 (Lomsadze *et al*., 2018). For Prokka, gene annotation used FASTA-format files as the ‘trusted’ CDS set (‘--protein’) if available, and tRNA and rRNA calling was turned off (‘--notrna’, ‘--norrna’). For Gene M arkS-2, genes were called using the online tool version (available at http://exon.gatech.edu/genemark/genemarks2.cgi) with default parameters.

### Pangenome analysis

Linear-genome pangenome analyses was conducted using Roary v3.13.0 (Page *et al*., 2015), Panaroo v1.2.10 (Tonkin-Hill *et al*., 2020) or PEPPAN v1.0.6 (Zhou, Charlesworth & Achtman, 2020) using gene annotations in GFF format provided by Prokka or GeneMarkS-2. All tools were run using default parameters, with the exception of Panaroo, which was run in moderate mode. ggCaller v1.3.3 was run on assemblies in FASTA-format in either sensitive, moderate, or strict modes. For simulated datasets, results were analysed using a custom script (compare_simulated_gene_pa.Rmd). For real datasets, gene frequency distributions were compared by generating histograms from gene presence/absence matrices in Rtab format from each workflow.

### Contig break analysis

Five manually annotated pneumococcal capsular polysaccharide synthesis operons from Bentley *et al*., (2006) were downloaded (Genbank accessions: CR931662.1, CR931663.1, CR931664.1, CR931665.1, CR931666.1). To fragment the operons, a single contig break was generated randomly in each manually annotated CDS using a custom script (fragment_at_gene.py). Gene predictions from each operon were compared to ground-truth gene sequences using a custom script (gene_recall.py). This script matches the 3’ ends between ground-truth and predicted genes to determine the number of correctly predicted complete sequences, and calculates the total proportion of ground-truth CDSs covered by gene predictions.

### Gene start/stop site comparison

Amino acid sequences for proteins within Pbp1a, Pbp2b, PsrP and PspA COGs were extracted from ggCaller and Prokka + Panaroo analyses of 616 *S. pneumoniae* genome sequences from Croucher *et al*., (2015). Sequences were aligned to reference protein sequences from Spn23F (Croucher *et al*., 2009) using Mafft v7.310 (Katoh *et al*., 2002). A custom script was used to identify soft clipping at the start and end of alignments compared to Spn23F sequences (gene_end_comparison.py). This script was also used to conducted all-by-all pairwise alignments within each COG to calculate average amino acid identity; the proportion of matching amino acids over the gapped alignment length (Doolittle, 1981; Raghava & Barton, 2006).

### Pangenome-wide association studies

616 *S. pneumoniae* genomes and their associated AMR MIC data were downloaded from Croucher *et al*., (2015). Genomes for which MIC data was available for tetracycline and erythromycin were extracted and analysed as separate datasets for each antibiotic. Isolates were labelled as either susceptible or resistant based on MIC cut-offs; 325/616 genomes had tetracycline MIC data and 36 isolates were labelled as resistant (MIC ≥8 μg/ml (Ousmane, Diallo & Ouedraogo, 2018)), 604/616 had erythromycin MIC data and 122 were labelled as resistant (MIC ≥1 μg/ml (Zhou *et al*., 2012)). Unitigs were identified in respective datasets using unitig-caller v1.2.1 (Lees *et al*., 2020). ggCaller was used to generate core-genome neighbour joining trees which were then midpoint-rooted. Unitigs and neighbour-joining trees were used to train mixed effects models with pyseer v1.3.10 (Lees *et al*., 2018) for respective datasets. Significant unitigs were identified using thresholds calculated by a built-in pyseer script (count_patterns.py); 2.42e^-07^ and 1.96e^-07^ for tetracycline and erythromycin respectively. All significant unitigs were mapped via exact alignment to Spn23F using the built-in pyseer annotation function (annotate_hits_pyseer.py) and to ggCaller-annotated DBGs using query mode in exact mapping (‘--query-id 1.0’) for respective datasets.

Mappings to Spn23F were visualised in Phandango (Hadfield *et al*., 2018). Mappings to the ggCaller graphs were analysed using a custom script (count_annotations.py), which determines the coverage of gene annotations by significant hits. Genes missing annotations were marked as ‘hypothetical’, whilst those annotated as transposons, insertion sequences, integrases or conjugative elements were marked as ‘MGE’ (mobile genetic element). To identify the genes with the greatest coverage of significant unitigs, genes with the same annotation were binned together, and the ratio of the total number of mapped queries to the total number of basepairs within each bin was calculated. This statistic is similar to fragments per kilo-basepair of transcript used in differential expression analysis (Zhao *et al*., 2021).

### Computational benchmarking

Genomes in FASTA-format from either the Global Pneumococcal Sequencing project (Gladstone *et al*., 2019) or Massachusetts dataset from Croucher *et al*., (2015) were randomly sampled using the ‘shuf’ bash command. Files were incrementally added to the subsample to increase dataset sizes. The same subsampled FASTA files were used for comparison of all workflows. All workflows were run with 16 threads on a server with 768 Gb memory and 2×20 core Intel Xeon Gold CPUs.

### Data access

The datasets generated and/or analysed in this manuscript are available in the ggCaller manuscript repository, https://github.com/samhorsfield96/ggCaller_manuscript. Source code for ggCaller can be accessed here: https://github.com/samhorsfield96/ggCaller. Documentation is available from readthedocs: https://ggcaller.readthedocs.io/en/latest/

## Acknowledgements

We thank the Bacterial Evolutionary Epidemiology group and the Pathogen Informatics and Modelling group at EMBL-EBI for their helpful comments during the development of ggCaller. In particular, we thank Dr Leonid Chindelevitch and Professor Nicholas Grassly at Imperial College London, and Professor Simon Frost at London School of Hygiene & Tropical Medicine for their support and advice.

## Authors’ contributions

STH, NJC and JAL conceived the project. STH developed the method and software and performed the data analysis. NJC and JAL and supervised the work and aided in interpretation of results. STH wrote the paper with contributions from all authors. All authors read and approved the final manuscript.

