## Supplemental materials for "Accurate and fast graph-based pangenome annotation and clustering with ggCaller"

### Supplementary figures

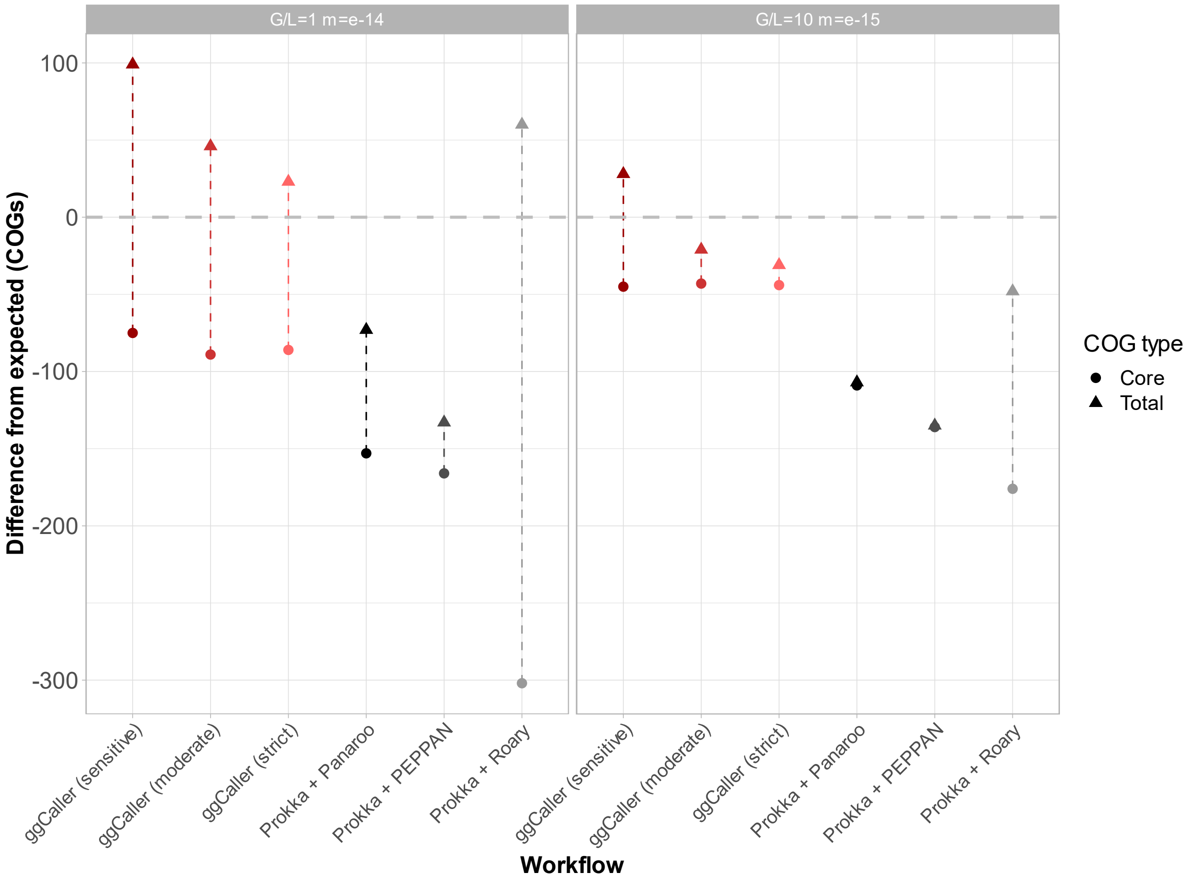

***Supplementary Figure 1: Comparison of estimated total pangenome and core genome sizes across simulated populations for simulations not included in Figure 3.*** Difference from expected refers to the over or underestimation of the number of COGs from the ground truth in the simulation (grey dotted line). G/L refers to the gene gain to gene loss ratio, m refers to the mutation rate per site per gene.

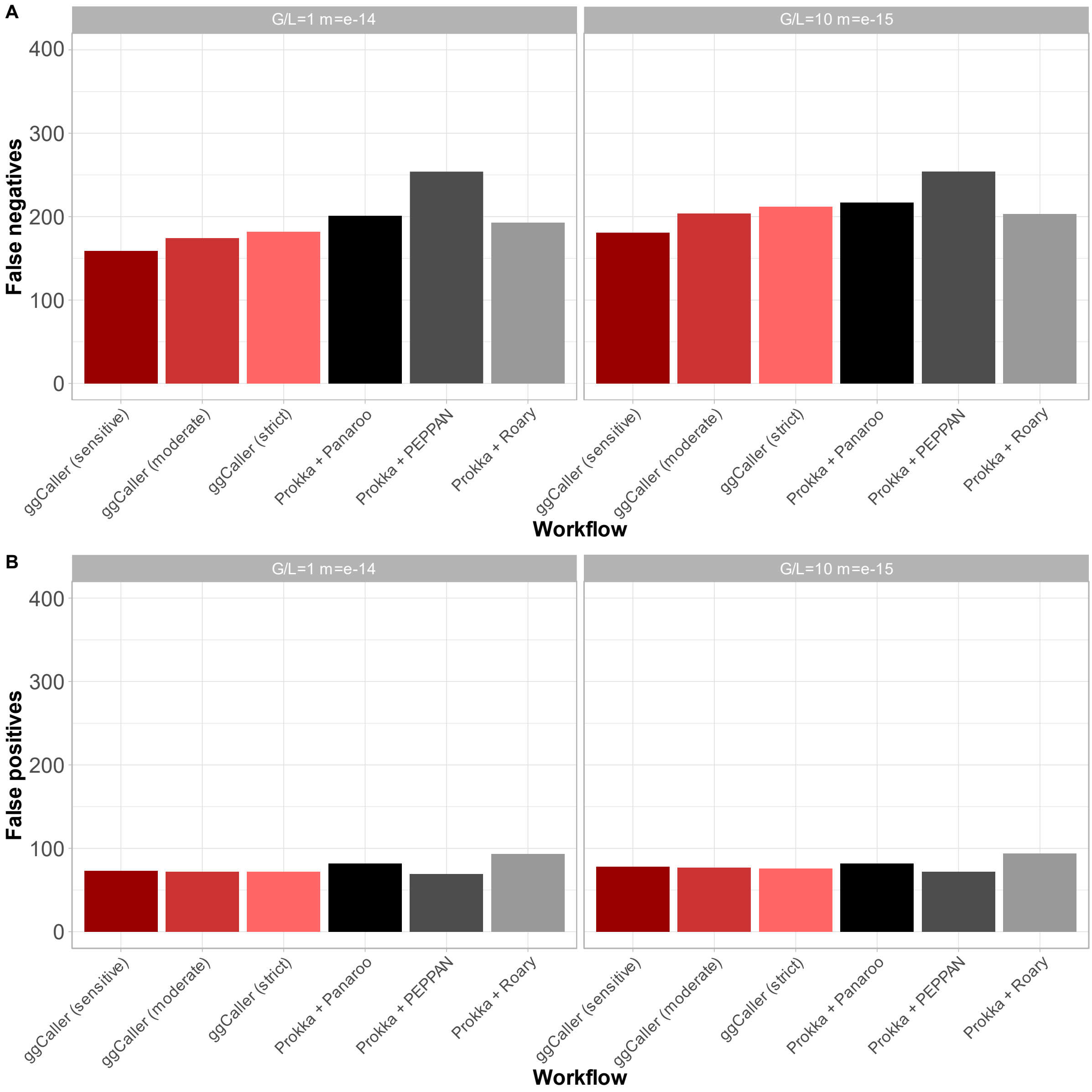

***Supplementary Figure 2: Comparison of COG annotation accuracy across simulated populations for simulations not included in Figure 4.*** Panels show false negatives (**A**) and false positives (**B**). False positives are COGs that were called by a workflow but were not present in the ground-truth set. False negatives are COGs that were present in the ground-truth set but not called by a workflow.

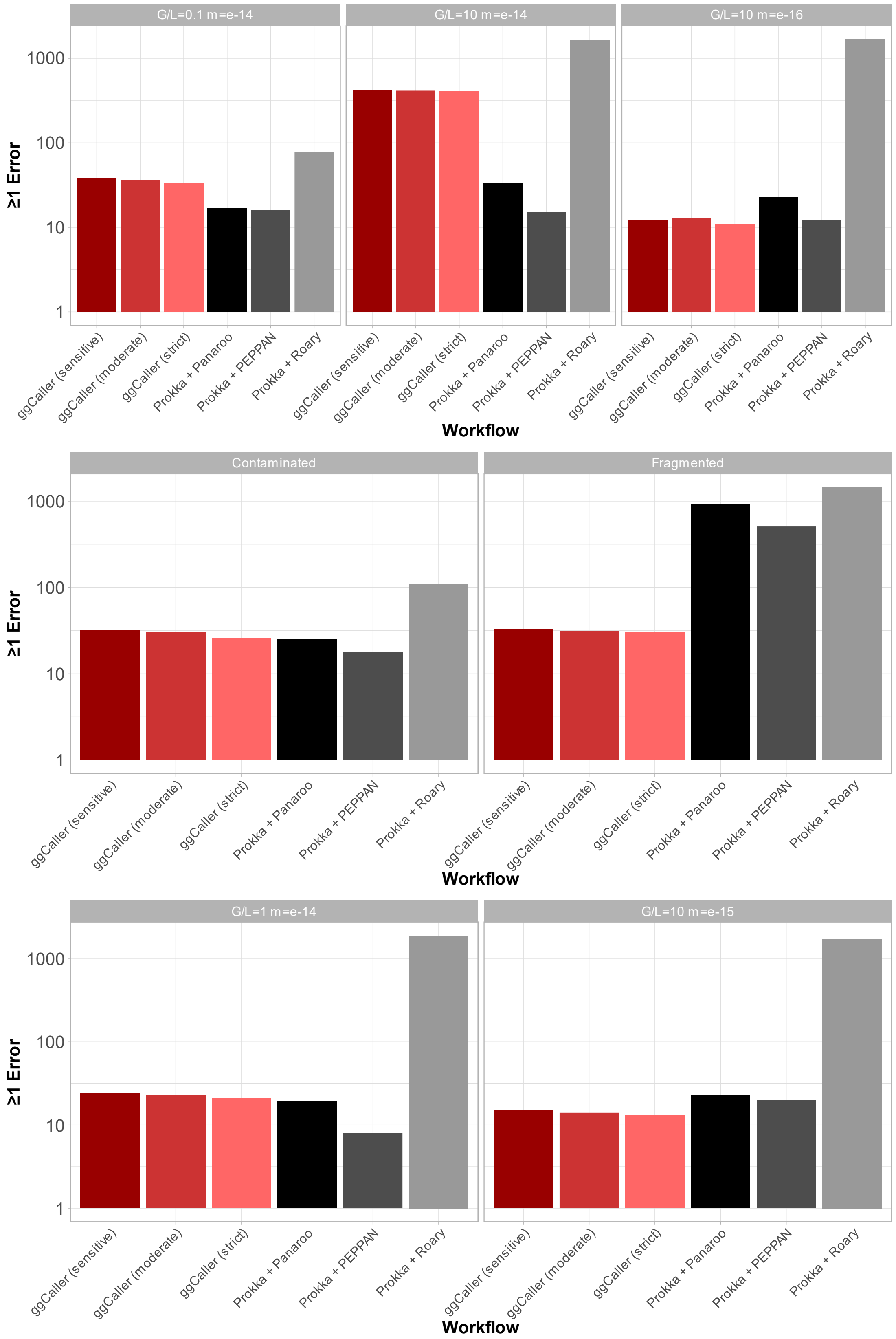

***Supplementary Figure 3: Comparison of number of COGs that contain at least one prediction error.*** For a COG to have a prediction error, it must contain an incorrectly predicted gene in one of the individual genomes. This can either be a gene called by a workflow which is missing in the ground truth (false positive), or a gene that is missed by a workflow present in the ground truth (false negative).

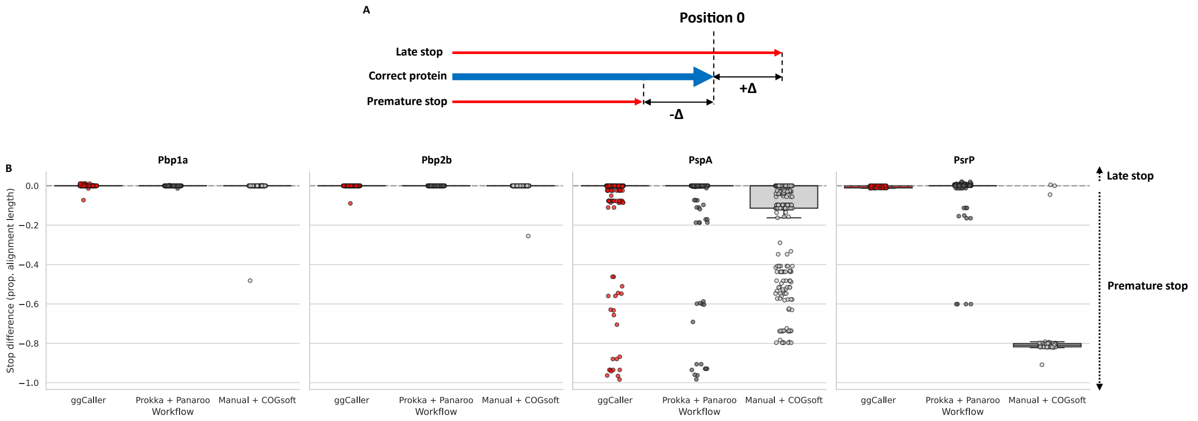

***Supplementary Figure 4: Comparison of within-COG stop site soft-clipping across ggCaller, Prokka + Panaroo and the original Massachusetts dataset.*** Description of stop site soft clipping (**A**). Comparison of stop site soft-clipping protein sequences of pbp1a, pbp2a, pspA and psrP based on alignment with the manually annotated reference in SPN23F (**B**).

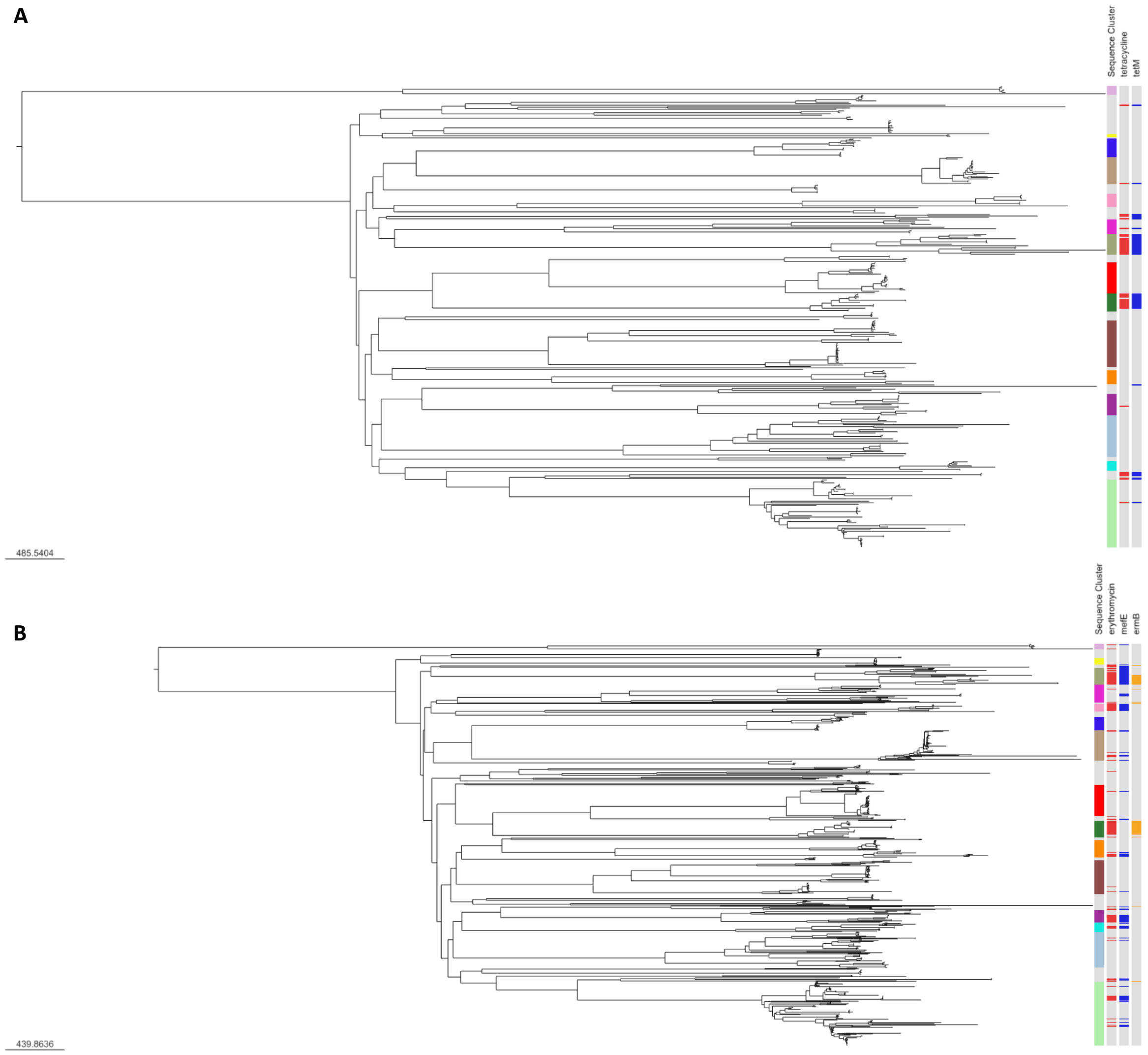

***Supplementary Figure 5: Core genome neighbour-joining tree generated by ggCaller annotated with resistance phenotype and AMR gene presence in S. pneumoniae.*** Trees were built from (A) 325 isolates with tetracycline MIC data and (B) 604 isolates with erythromycin data (Croucher et al., 2015). Trees were displayed using Microreact (Argimón et al., 2016). Key for core SNP-distance to branch length shown in bottom left corner of each panel. Blocks to right of trees describe sequence clusters assigned in Croucher *et al.*, drug resistance phenotype and AMR gene presence identified by ggCaller.

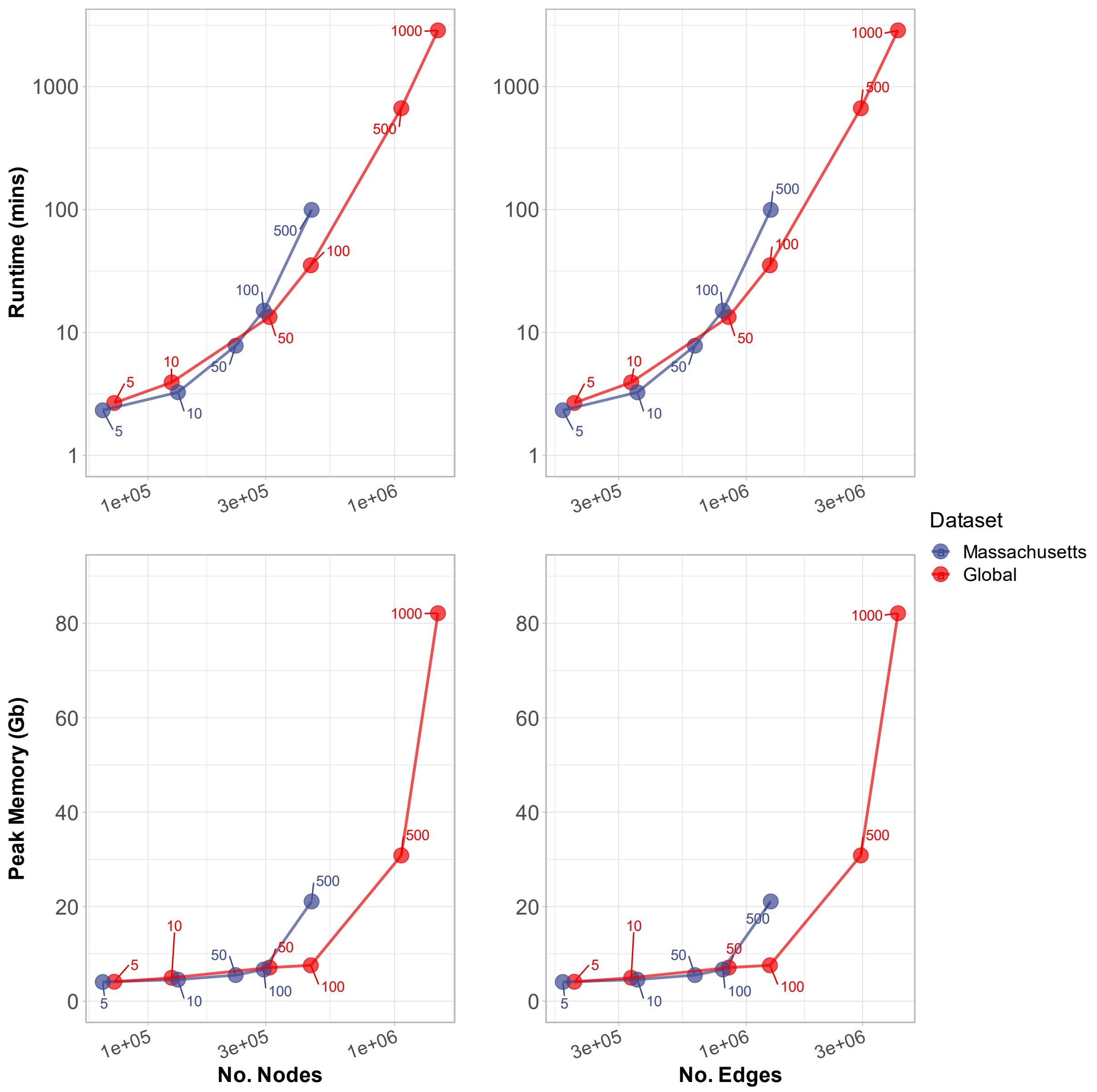

***Supplementary Figure 6: Comparison of de Bruijn Graph complexity and ggCaller computational performance.*** Nodes refer to unitigs within a de Bruijn Graph, edges refer to connections between nodes. Point labels indicate the number of isolate genomes included in analysis. All analyses were run with 16 threads.

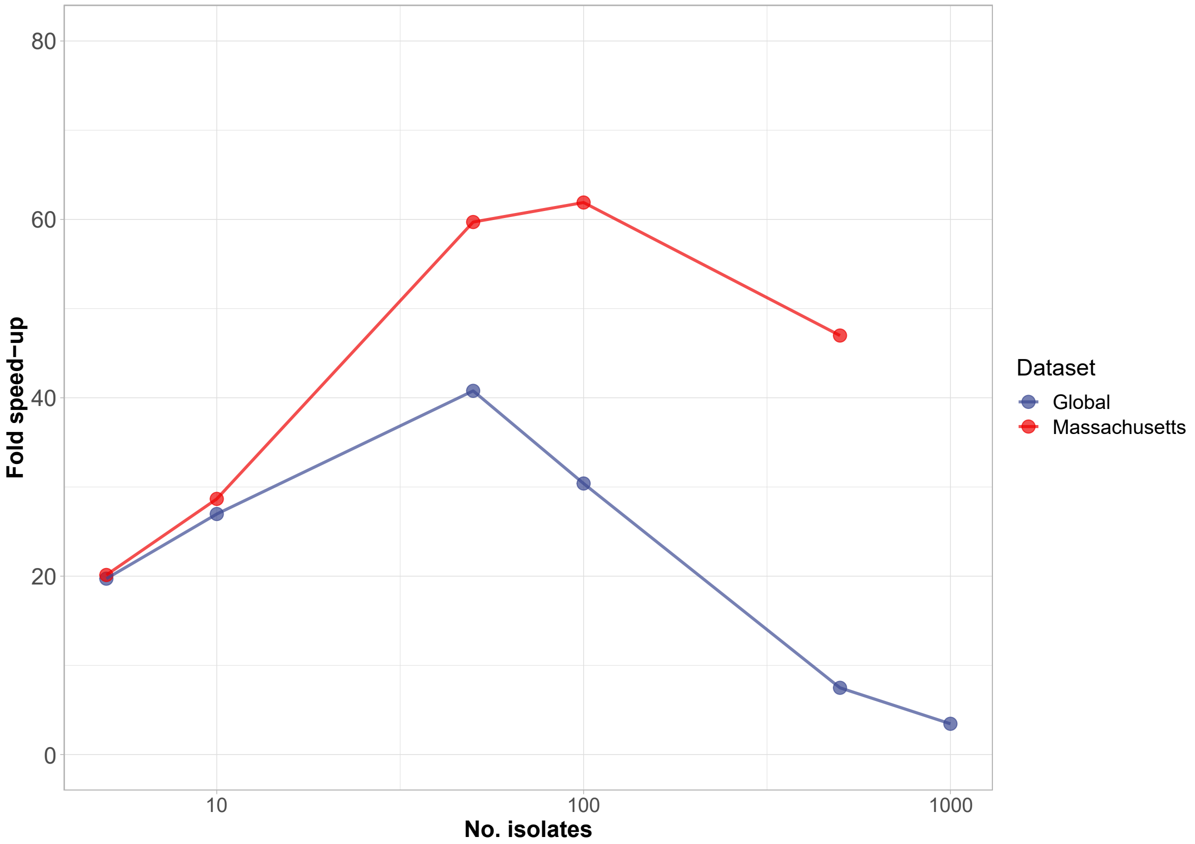

***Supplementary Figure 7: Fold speed-up of ggCaller over Prokka + Panaroo for Global and Massachusetts datasets of Streptococcus pneumoniae for increasing dataset size.*** Fold speed-up was calculated by dividing the runtime of Prokka + Panaroo by that of ggCaller for runs with identical datasets. Both workflows were run with 16 threads.

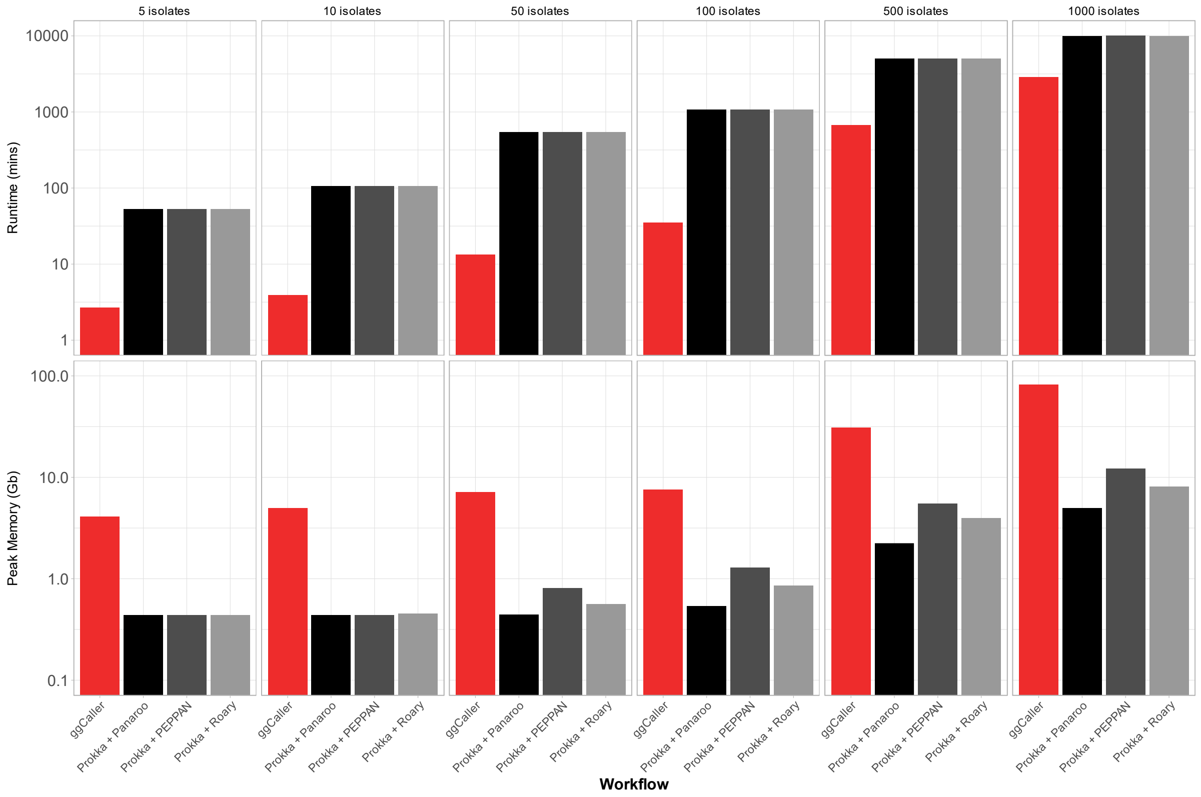

***Supplementary Figure 8: Computational performance comparison between ggCaller and pangenome analysis workflows on global Streptococcus pneumoniae dataset.*** All tools were run using 16 threads.

### Supplementary tables

***Supplementary Table 1: Simulation parameters using infinitely many genes model.*** Gene gain and loss rates are measured in per-generation per genome. Gene mutation rate is measured in per-generation per-nucleotide. No. genes is the number of genes simulated (i.e. those in accessory genome), remainder are left as core genes. No. isolates determines how many genomes are simulated. Effective population size determines the population size in terms of the number of gene copies.

| Name | Gene gain rate | Gene loss rate | Gene mutation rate | No. Genes | No. Isolates | Effective population size | Additional |
|---|---|---|---|---|---|---|---|
| G/L=0.1 m=e-14 | $1e^{-13}$ | $1e^{-12}$ | $1e^{-14}$ | 1000 | 100 | $10^6$ | NA |
| G/L=1 m=e-14 | $1e^{-12}$ | $1e^{-12}$ | $1e^{-14}$ | 1000 | 100 | $10^6$ | NA |
| G/L=10 m=e-14 | $1e^{-12}$ | $1e^{-13}$ | $1e^{-14}$ | 1000 | 100 | $10^6$ | NA |
| G/L=10 m=e-15 | $1e^{-12}$ | $1e^{-13}$ | $1e^{-15}$ | 1000 | 100 | $10^6$ | NA |
| G/L=10 m=e-16 | $1e^{-12}$ | $1e^{-13}$ | $1e^{-16}$ | 1000 | 100 | $10^6$ | NA |
| Contaminated | $1e^{-12}$ | $1e^{-13}$ | $1e^{-14}$ | 1000 | 100 | $10^6$ | 10kb fragmented Staphylococcus Epidermidis added per genome (insert_random_genome_fragments.py) |
| Fragmented | $1e^{-12}$ | $1e^{-13}$ | $1e^{-14}$ | 1000 | 100 | $10^6$ | Genomes fragmented (fragment_fasta.py) |
